# Double Reduction in Allotetraploid Peanut and the Role of Chromosomal Imbalance in Unexpected Linkage Map Artifacts

**DOI:** 10.64898/2026.02.12.704920

**Authors:** Samuele Lamon, Peter M. Bourke, Brian L. Abernathy, João F. dos Santos, Ignácio J. de Godoy, Soraya C.M. Leal-Bertioli, David J. Bertioli

## Abstract

Polyploidization in peanut (*Arachis hypogaea* L.) provided evolutionary advantages by increasing heterosis, the response to selection, and enhancing adaptability. However, it also caused a genetic bottleneck by isolating cultivated peanut from its wild diploid relatives. Mechanisms such as homoeologous exchange can partially restore genetic diversity by generating new allelic combinations. Double reduction is a rare segregation pattern restricted to polyploids, in which a single-dosage locus yields duplex gametes. It requires multivalent formation and crossing over between non-sister chromatids, both of which are associated with homoeologous exchange. Although peanut mainly exhibits disomic pairing, occasional multivalents theoretically allow low-frequency double reduction. To estimate double reduction and examine its relationship with genetic instability, a high-density phased linkage map was constructed using a backcross population from a cross between a neoallotetraploid [*A. magna* K 30097 x *A. stenosperma* V 15076]^4x^ (MagSten) and cultivated peanut. The final map included 9,717 SNP markers with an average spacing of 0.22 centiMorgans. Some progenies showed unbalanced genomic compositions, creating artifacts in linkage analysis. Removing these progenies improved the map and suggested a common origin for artifacts previously observed in other linkage maps, revealing a novel aspect of mapping in allotetraploid peanut. Analysis of the phased map revealed double reduction in 12% of progenies. Notably, one event produced a genomic composition consistent with theoretical predictions, supporting the expectation that double reduction causes unbalanced genomes in allopolyploids. These results indicate that double reduction is a low but significant frequency genetic phenomenon in the segmental allotetraploid peanut, contributing to the genetic instability and evolutionary dynamics of this and likely other allopolyploid genomes.

**Article Summary:** This study investigated double reduction, a rare genetic event in segmental allopolyploid peanut, which can create unbalanced genomic compositions and affect genetic diversity. We generated a backcross population using neoallotetraploid and cultivated peanuts, then constructed a high-density phased linkage map. Analysis revealed unbalanced genomic compositions in some progenies caused by homoeologous exchanges, which reduced map quality. Double reduction was estimated to occur in approximately 12% of progenies, aligning with theoretical expectations for genomic imbalance. These results demonstrate that double reduction contributes to genetic instability, inheritance patterns, and genome evolution in allopolyploid organisms such as peanut.

## Introduction

Peanut (*Arachis hypogaea* L.) is an allotetraploid species (2n = 4x = 40; AABB-type genome) that originated less than 10,000 years ago through a spontaneous hybridization event involving two diploid wild *Arachis* species, *A. duranensis* Krapov. & W.C. Greg. (A-type genome) and *A. ipaënsis* Krapov. & W.C. Greg. (B-type genome). The hybridization of these two wild species was likely facilitated by humans, who introduced *A. ipaënsis* into the growing habitat of *A. duranensis* (Bertioli et al. 2016; Bertioli et al. 2019; de Blas et al. 2025).

Either through unreduced gametes or through hybridization followed by whole genome duplication (Garcia et al. 2020), cultivated peanut became tetraploid and genetically isolated from its wild diploid *Arachis* relatives. This event led to a significant genetic bottleneck in the peanut population and resulted in a drastic reduction of genetic diversity within the peanut gene pool (Kochert et al. 1996; Seijo et al. 2007; Bertioli et al. 2011; Moretzsohn et al. 2013; Bertioli et al. 2019).

However, the merging of two distinct diploid genomes in peanuts triggered a ‘genetic shock,’ resulting in genetic instability, large-scale deletions and inversions (McClintock 1984; Bertioli et al. 2019; Mason and Wendel 2020). As a result, peanuts evolved morphological diversity, eventually becoming a globally recognized crop, aided by an increased response to selection compared to their diploid relatives (Lamon et al. 2025a). Nowadays, peanuts are classified into two subspecies: *hypogaea* and *fastigiata*, and six botanical varieties - *hypogaea* and *hirsuta* (within the *hypogaea* subspecies), and *fastigiata*, *vulgaris*, *aequatoriana*, and *peruviana* (in the *fastigiata* subspecies). Additionally, peanut encompasses four major cultivated market types: Runner, Spanish, Valencia, and Virginia, and thousands of landraces and cultivars, each one with its own distinct characteristics (Bertioli et al. 2011, 2019).

Similar destabilizing mechanisms, including homoeologous exchange and its associated unbalanced chromosomal compositions, such as BBBB, ABBB, AAAB, and AAAA configurations, are still observed in modern cultivated peanut (Lamon et al. 2025b). Understanding the drivers of genetic instability may offer deeper insights into the mechanisms underlying increased genetic and morphological diversity in peanut, as well as its enhanced response to selection.

Double reduction (DR) is that circumstance in polyploid organisms where an individual carrying a locus in a single dosage produces duplex gametes for that same locus during meiosis (Darlington 1929). DR increases homozygosity, diminishes the frequency of deleterious alleles in a population over time through negative selection and has the potential to increase rare advantageous alleles in plant material if combined with marker-assisted selection (Butruille and Boiteux 2000; Bourke et al. 2015). In order for DR to manifest, three main events need to occur: multivalent pairing of chromosomes during prophase I, followed by crossing-over between non-sister chromatids, and a subsequent non-disjunctional chromosomal separation pattern, which implies that first and second meiotic divisions are reductional for the considered loci (Mather 1936; Ronfort et al. 1998; De Silva et al. 2005; Huang et al. 2019). The first two requirements, multivalent formation and crossing-over between non-sister chromatids, are commonly associated with homoeologous exchange (Mason and Wendel 2020).

Polysomic inheritance and multiple chromosomal pairing are typical of autopolyploid organisms since extremely high similarity is shared amongst different sets of chromosomes. In autopolyploids, ploidy increase takes place within a single species, often in a sole individual. Conversely, allopolyploids are formed by the hybridization of two different species and are also generally characterized by disomic inheritance (resulting from chromosome pairing within but not between subgenomes). In 2002, Ramsey and Schemske estimated that on average the percentage occurrence of multivalents in autopolyploid and allopolyploid plants are 28.8% and 8.0%, respectively. Even though multivalents are rarer in allopolyploids than autopolyploids, they arguably have a much higher biological significance in such species, given that recombination has a higher impact if genomes are more divergent (McGrath and Lynch 2012).

Detecting DR is a complex process that often requires a focus on specific markers and cross-combination approaches. Similarly, accurately estimating DR rate can be challenging, frequently relying on computer simulations (Tai 1982; Haynes and Douches 1993; Bourke et al. 2015).

Percentages of DR have been mainly detected and estimated in autopolyploid plants e.g. purple loosestrife (*Lythrum salicaria* L.) (Fisher 1943; Fisher and Mather 1943), potato (*Solanum tuberosum* L.) (Haynes and Douches 1993; Bourke et al. 2015), great yellowcress (*Rorippa amphibia* (L.) Besser) and yellow fieldcress (*R. sylvestris* (L.) Besser) (Stift et al. 2008), purple yam (*Dioscorea alata* L.) (Nemorin et al. 2012), clementine (*Citrus clementina* Hort. ex Tan.) (Aleza et al. 2016), hybrid tea rose (*Rosa hybrida* L.) (Bourke et al. 2017), chrysanthemum (*Chrysanthemum* × *morifolium* (Ramat.) Hemsl.) (Bourke et al. 2021) and much more rarely in animals such as rainbow trout (*Oncorhynchus mykiss* Walbaum) (Allendorf and Danzmann 1997) or Pacific oysters (*Crassostrea gigas* (Thunberg)) (Curole and Hedgecock 2005).

In contrast, the detection and quantification of significant DR frequencies in allopolyploid organisms have been much more challenging. To our knowledge, DR has not been conclusively demonstrated in any confirmed allopolyploid species. Ahmed et al. (2020) estimated DR levels in the ‘Giant Key’ lime, which is likely a doubled diploid derivative of the ‘Mexican’ lime (*C. aurantifolia* (Christm.) Swingle). However, the precise polyploid origin of this species remains uncertain.

Peanut follows a third genetic model, called segmental allopolyploidy. As such, it exhibits mostly disomic paring, with occasional multivalent formations and homoeologous exchanges (Husted 1933, 1936; Stalker 1981; Leal-Bertioli et al. 2015a). These features make the occurrence of DR theoretically possible in peanut. However, due to the rarity of multivalents and high homozygosity in cultivated peanut, detecting and estimating DR remains challenging. To overcome this limitation, we generated a BC_1_ population using the neoallotetraploid [*A. magna* K 30097 × *A. stenosperma* V 15076]^4x^ (MagSten) (Fávero et al. 2015) as the donor parent and ‘IAC OL4’ (*A. hypogaea*) (de Godoy et al. 2014) as the recurrent parent (Moretzsohn et al. 2023). This cross generated sufficient polymorphism to track individual alleles and test for the presence of DR.

In this study, we estimated DR occurrence, with direct evidence of an event associated with an AAAB genomic composition. Both theoretical considerations and our findings indicate that, because DR is associated with homoeologous exchange, its occurrence in an allopolyploid species such as peanut frequently results in unbalanced genomic compositions in the progenies (Figure 1). To enable accurate estimation of DR, we constructed a high-density, phased genetic linkage map, which facilitated detailed analysis of allele inheritance patterns. Additionally, we identified double linkage group (LG) artifacts in Marey plots, which resulted from large-scale unbalanced genomic compositions caused by homoeologous exchange. Resolving these artifacts improved the resolution of the genetic map and demonstrated how genetic instability can distort linkage analysis in segmental allopolyploid species such as peanut.

**Figure 1:**
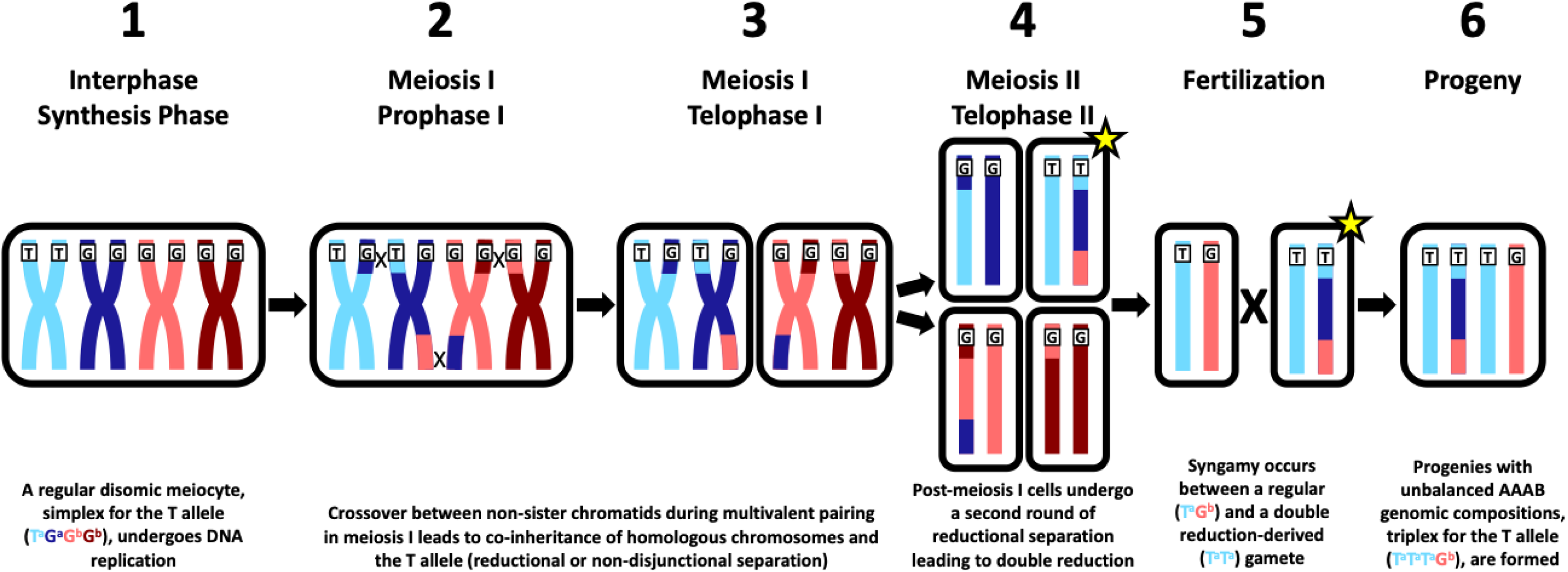
Mechanism of double reduction (DR) in segmental allotetraploid peanut. Letters “G” and “T” represent two different alleles. Blue (both light and dark) and red (both light and dark) colors, as well as superscripts “a” and “b”, indicate the two peanut subgenomes, A and B, respectively. 1) During the synthesis phase of interphase, a disomic meiocyte with a simplex genotype for the “T” allele (T^a^G^a^G^b^G^b^) undergoes DNA replication, forming sister chromatids. Although this heterozygous configuration is rare in peanut due to high subgenome homozygosity, the backcross population structure used in this study provides sufficient polymorphism for such configurations to occur (see Figure 2). 2) In prophase I, due to the segmental allopolyploid nature of peanut, both homologous and, occasionally, homoeologous chromosomes can pair, forming multivalents. For homologous and homoeologous chromosomes to pair, crossing over needs to occur between non-sister chromatids (homoeologous exchange). 3) Multivalent pairing and homoeologous exchange allow two homologous chromosomes carrying the “T” allele to be inherited together into the same daughter cell (reductional or non-disjunctional separation), which is not possible under strict disomic inheritance. 4) During meiosis II, a second round of chromatid segregation occurs. The two “T” alleles may again segregate into the same gamete (second round of reductional or non-disjunctional separation), leading to the formation of a gamete with two copies of the “T” allele, which in other words is DR (highlighted by yellow star). 5) During fertilization, the DR-derived gamete will very likely pair with a regular gamete resulting from disomic inheritance (T^a^T^a^G^b^G^b^ generates T^a^G^b^ gametes), given that inheritance is mainly disomic in peanut. 6) Progenies resulting from DR will likely have unbalanced AAAB genomic compositions (T^a^T^a^T^a^G^b^) at the considered locus. This demonstrates how DR increases homozygosity but also generates unbalanced genomic compositions at the considered locus in segmental allotetraploid species such as peanut.

## Materials and Methods

### Plant material

A BC_1_ population was developed using the neoallotetraploid [*A. magna* K 30097 × *A. stenosperma* V 15076]^4x^ (MagSten) (Fávero et al. 2015) as the donor parent and ‘IAC OL4’ (*A. hypogaea*) (de Godoy et al. 2014) as the recurrent parent (Moretzsohn et al. 2023). A total of 74 BC_1_ progenies, numbered from’Ind 1’ to‘Ind 74,’ were included in the study. Additionally, BC_1_F_2_ progenies were derived by selfing from Ind 27, 40, and 41. These BC_1_F_2_ progenies were labeled based on single-seed descent identification. For example, Ind 40.2 indicates the second progeny of the BC_1_ plant Ind 40.

### Genotyping and sequencing

DNA was extracted from young leaves of BC_1_ progenies, selected BC_1_F_2_ progenies, and ‘IAC OL4’ and MagSten parentals. Genotyping was performed using the Axiom *Arachis* 48K SNP array v2 (Affymetrix, Santa Clara, CA, USA) (Pandey et al. 2017; Korani et al. 2019). In addition, four BC_1_ progenies (Ind 33, 39, 47, and 62) and two BC_1_F_2_ progenies (Ind 40.3 and 40.4) underwent whole-genome sequencing (WGS) at 30× coverage using PCR-free Illumina NovaSeq PE150 technology (Novogene Co., Sacramento, CA, USA).

### Genotypic data scoring and linkage mapping

Raw.CEL files from the Axiom *Arachis* 48K SNP array v2 were processed using the Axiom Analysis Suite 2022 (Thermo Fisher Scientific, Waltham, MA, USA) to generate a summarized signal intensity file, which was then imported into RStudio for further analysis (Posit Team 2025). SNP markers for tetraploid samples were scored on a tetraploid scale (0 to 4) using the ‘fitPoly’ package (Voorrips, Gort and Vosman 2011; Voorrips and Gort 2018), with ‘IAC OL4’ and MagSten given as parents of the BC_1_ population, as the F_1_ hybrid genotypic data were unavailable (see Supporting Information, Table S1).

Markers filtering and linkage mapping were performed using ‘polymapR’ (Bourke et al. 2018). Initially, markers were filtered for non-missing values in the parents, retaining 25,315 markers (Figure S1A). Subsequently, non-segregating markers were removed and dosages were converted to basic segregating types, resulting in a final set of 11,565 markers (Figure S1B). Next, markers and individuals of the BC_1_ population were filtered for missing values, applying a cutoff rate of 0.25 for both. The allele dosage matrix was then filtered for duplicated individuals (cutoff value 0.95), identifying four pairs of duplicated individuals and four individuals resulting from self-pollination in ‘IAC OL4’. After merging these cases, 66 individuals remained for linkage mapping. Finally, the allele dosage matrix was filtered for duplicated markers, retaining 5,225 unique markers (Figure S1C).

Most markers were initially scored by ‘fitPoly’ as simplex x nulliplex and nulliplex x simplex types (Figure S1, B and C). However, because cultivated peanut is characterized by extremely high subgenome homozygosity, simplex x nulliplex markers are expected to be very rare. As MagSten, was entered in place of the F_1_ hybrid the pollen parent for genotype calling, it is more accurate to interpret these simplex x nulliplex markers as actually representing duplex x simplex segregation types (Figure 2A). For further mapping purposes, all markers erroneously called as simplex x nulliplex were re-labelled as nulliplex x simplex.

**Figure 2:**
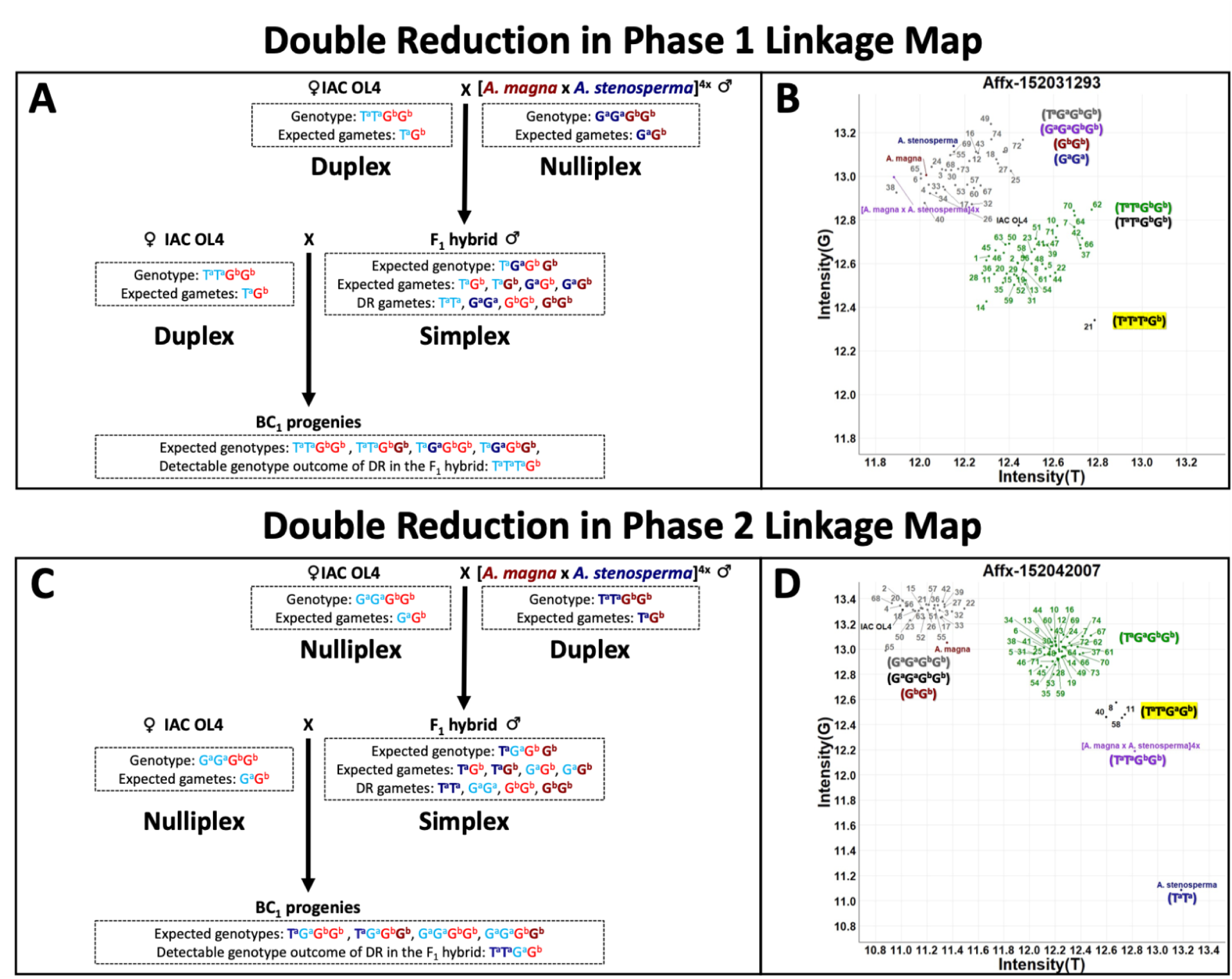
Double reduction (DR) occurrence in segmental allotetraploid peanut. Diagrams show DR events in the ‘IAC OL4’ × F_1_ hybrid BC_1_ population. Panels A and B illustrate linkage map phase 1; panels C and D show phase 2. “G” and “T” indicate alleles, with superscripts “a” and “b” for peanut subgenomes. In panels A and C, colors indicate *Arachis* genome-species combinations: ‘IAC OL4’ (light blue/red for a/b), *A. magna* (b; dark red), and *A. stenosperma* (a; dark blue). In panels B and D, numbers label BC_1_ progenies, with y-axis showing “G” allele intensity and x-axis “T” allele intensity.

Panel A: If ‘IAC OL4’ is duplex (T^a^T^a^G^b^G^b^) and MagSten nulliplex for “T” (G^a^G^a^G^b^G^b^), the F_1_ hybrid is simplex (T^a^G^a^G^b^G^b^), producing T^a^G^b^ or G^a^G^b^ gametes under disomic inheritance, while ‘IAC OL4’ produces only T^a^G^b^. DR in the F_1_ hybrid can generate T^a^T^a^ gametes, resulting in unbalanced AAAB triplex compositions (T^a^T^a^T^a^G^b^), distinguishable from simplex/duplex scores. Panel B: SNP intensities (Affx-152031293) reflect panel A, with color-coded genotype-genome-allele combinations; AAAB DR-derived BC_1_ progeny (T^a^T^a^T^a^G^b^) is highlighted in yellow.

Panel C: If ‘IAC OL4’ is nulliplex (G^a^G^a^G^b^G^b^) and MagSten duplex for “T” (T^a^T^a^G^b^G^b^), the F_1_ hybrid is simplex (T^a^G^a^G^b^G^b^), producing T^a^G^b^ or G^a^G^b^ gametes under disomic inheritance, while ‘IAC OL4’ yields only G^a^G^b^. DR in the F_1_ hybrid can generate T^a^T^a^ gametes, producing unbalanced AAAB duplex compositions (T^a^T^a^G^a^G^b^), distinguishable from nulliplex/simplex scores.

Panel D: SNP intensities (Affx-152042007) reflect panel C, with color-coded genotype-genome-allele combinations; AAAB DR-derived BC_1_ progenies (T^a^T^a^G^a^G^b^) are highlighted in yellow.

Markers were aligned against A and B subgenomes of the ‘Tifrunner’ (*A. hypogaea*) reference genome (version gnm2) to determine their physical positions (Bertioli et al. 2019). The alignment process was conducted in the Unix environment.

Pairwise recombination frequency, LOD scores, and linkage phases were calculated for markers using the ‘polymapR’ linkage() function with ploidy equal 2, with an expectation of 20 LGs. Marker clusters were generated over a range of LOD thresholds, with a minimum cluster size of 11 markers; at a threshold of 11, unique markers were clustered into 20 groups. Concordance of marker positions across the two reference assemblies, along with the application of the phase_SN_diploid() function, was used to phase markers between the A and B subgenomes. Inter-LG linkage signal was then assessed and duplicated markers were reintegrated, yielding a total of 9,717 markers (see Supporting Information, Table S2).

The classification of markers during genotype calling into duplex × simplex (re-labelled as nulliplex x simplex from mapping purposes) and nulliplex × simplex effectively phased markers between the two parental lines, with duplex × simplex corresponding to phase 1 (‘IAC OL4’) and nulliplex × simplex corresponding to phase 2 (F_1_ hybrid). A very high concordance was observed between “genotyping calling phasing” and linkage map phasing (Figure S2).

Summary statistics were calculated for the integrated map, and Marey plots were generated to illustrate the relationship between genetic distances and the physical positions on the ‘Tifrunner’ reference genome, with genetic positions reoriented where necessary to match the physical map; pericentromeric regions were highlighted. Out of 9,717 markers, 1,628 were excluded from the plots because their corresponding subgenomes lacked ‘Tifrunner’ physical positions. The plots revealed unexpected double LG artifacts in LGs 4 and 14, with the artifact in LG14 being so pronounced that it resulted in the formation of two distinct LGs.

### Interspersed genotyping patterns and Marey plots resolution

During linkage map construction, we observed pronounced mapping artifacts in some LGs, including the appearance of doubled lines and even an entire doubled LG in Marey plots. These artifacts suggested that a subset of progenies might violate common assumptions (such as balanced tetraploid inheritance) used for linkage analysis.

After initial map refinement by removing markers with missing alleles or lacking physical alignment information, the integrated map was divided into two phases. Examination of the phased maps—particularly phase 2—revealed extensive interspersed genotyping patterns, characterized by rapid switches in inferred allele dosage between consecutive markers ordered by physical position. Such interspersed genotyping patterns are indicative of unbalanced polyploid states (e.g. AAAB or ABBB) that produce allele dosage states incompatible with the genotyping model assumptions and have previously been described by Lamon et al. (2025a). In that study, unbalanced genotypic compositions could not be resolved due to limited dosage discrimination but were partially mitigated by scoring genotypes on a tetraploid scale using ‘fitPoly’ and restricting analyses to markers with clear A- and B-subgenome differentiation. In the present study, however, resolving interspersed genotyping proved more challenging.

Although BC₁ progenies were scored on a tetraploid scale, the mapping population comprised four distinct subgenomic sources: A and B subgenomes from the cultivated parent (‘IAC OL4’) and A and B subgenomes from the wild parents (*A. stenosperma* and *A. magna*). In phase 1, A- and B-subgenome alleles were distinguishable in the cultivated parent but identical in the wild parent, whereas the opposite configuration occurred in phase 2 (Figure 2). This phase-dependent subgenomic symmetry reduced effective dosage resolution and likely contributed to interspersed genotyping patterns, similar to those previously reported, complicating the identification of progenies with unbalanced genomic compositions.

To systematically identify such cases, we developed an algorithm to detect regions of interspersed genotyping. Phased colormaps were first converted into binary matrices indicating whether allele dosage changed between consecutive markers. A sliding window of 11 markers (five upstream, the focal marker, and five downstream) was applied; if more than four dosage changes occurred within the window, the central marker was classified as part of an interspersed genotyping region. These regions were visualized as chromosome-level heatmaps, and the number of interspersed markers per chromosome was summarized using bar plots.

Interspersed genotyping was most pronounced on LG14 of phase 2, which was also the LG exhibiting strong double-LG artifacts in Marey plots. To test whether these artifacts were caused by progenies with abnormal genomic composition, we quantified interspersed genotyping on LG14 (phase 2) for each BC₁ progeny. This analysis identified 18 progenies with markedly elevated interspersed patterns (Figure S3A).

Linkage mapping was then repeated after excluding these progenies. This intervention resulted in the complete disappearance of the double LG artifacts, particularly on LGs 4 and 14, and restored coherent Marey plot structure. These results demonstrate that large-scale unbalanced genomic compositions—manifesting as interspersed genotyping patterns—were responsible for the observed double-LG artifacts, rather than errors in marker ordering or map construction.

### Estimation of DR

The integrated linkage map was refined by removing markers with missing alleles or lacking physical alignment information and subsequently divided into the two phases. Regions showing DR-type segregation were identified. While similar patterns could also result from tetrasomic recombination in the female parent during the backcross in phase 1 or from two consecutive rounds of tetrasomic recombination in the male parent in phase 2, DR provides the most parsimonious explanation for the observed patterns, especially in phase 2 (Figure 2).

To minimize the impact of genotyping errors, only regions where multiple consecutive markers displayed DR scores were considered as true DR events. This filtering step was performed manually through visual inspection of phased colormaps. Final DR inferences for both phases were visualized using heatmaps using ‘ggplot2’, providing a chromosome-level overview of DR event distribution.

### DR estimation using a hidden Markov model

To independently validate DR inference, we explored an alternative approach based on estimating posterior genotype probabilities using a multivalent-compatible Hidden Markov Model (Zheng et al. 2016). However, such an approach would require an integrated genetic linkage map, which is impossible to create due to lack of linkage between subgenomes (e.g. LG1 and LG11 are syntenic with chromosome 01 but cannot be combined into a single linkage map). To circumvent this difficulty, we used a physical-assembly guided approach to create a pseudo-integrated linkage map of ten groups via centromere alignment. Firstly, pericentromeric regions were identified visually from Marey maps (genetic vs physical marker positions) using consensus physical positions (Figure S4A). The median genetic position of markers within these centromeres was taken as the approximate genetic position of the centromeres, and for each chromosome, one of the maps was shifted upwards so that centromeres were genetically aligned (Figure S4B). A pseudo-integrated map was created by merging both maps and re-sorting markers on the new consensus genetic positions (Figure S4C, Figure S5). This was converted to a phased map by concatenating the phase information per chromosome across subgenomes into a “pseudo-autotetraploid” phased map (adding a dummy phase for parent 1, for which no segregating markers were present).

The pseudo-integrated phased map and marker genotypes were used to estimate posterior genotype (IBD) probabilities in ‘polyqtlR’ v.0.1.1 (Bourke et al. 2021) using the estimate_IBD() function with settings ploidy = 4 and bivalent_decoding = FALSE (i.e. allowing for the possibility of multivalent formation). These probabilities were subsequently re-estimated using the maxL_IBD() function using a range of genotyping error priors (ɛ = 0.01, 0.05, 0.1, 0.15, 0.2, 0.25, 0.3, 0.35). Maximum likelihood estimates for the genotyping error rates per individual per chromosome were visualized in a heatmap.

### Sequencing data analysis

To assess subgenome balance and evaluate genomic composition in selected progenies, we first identified a robust set of subgenome-diagnostic polymorphisms. This was done by using independent sources of short sequences aligned to a common reference genome: *in silico*–fragmented assemblies of *Arachis duranensis* (A genome) and *A. ipaënsis* (B genome), and Illumina WGS data from two independent sources. Using multiple sources ensured that subgenome-specific alleles were consistently identifiable and reliably recoverable from Illumina sequencing data in the progenies, following Lamon et al. (2025a). Illumina reads were quality-filtered using BBMap (v39.01) (Bushnell 2014) and, together with the *in silico*–fragmented assemblies, were aligned to the *A. ipaënsis* gnm1 reference genome using BWA (v0.7.17-r1188) (Li and Durbin 2009), which provided a common coordinate framework. Variant calling was performed using BCFtools (version 1.15.1) (Danecek et al. 2021) to identify positions that consistently differentiated the A and B subgenomes. AB-diagnostic variant positions were retained only if they met stringent criteria, including concordance between sequencing datasets, a minimum of 95% consensus among mapped reads, sufficient read depth, and clear allele differentiation between the A and B subgenomes. This procedure yielded a fixed set of high-confidence subgenome-specific polymorphisms.

Genomic composition in progenies was inferred by calculating normalized counts of subgenome-diagnostic alleles at the predefined AB-diagnostic positions. Illumina WGS reads from selected BC₁ and BC₁F₂ progenies (Ind 33, 39, 47, 62, 40.3, and 40.4) were then aligned to the same reference genome, and allele counts were extracted exclusively at these positions. This targeted interrogation allowed quantitative assessment of subgenome allele representation without *de novo* variant discovery. To normalize allele counts and account for potential mapping and coverage biases, synthetic tetraploid controls were generated by combining HudsonAlpha Illumina WGS reads from *A. duranensis* and *A. ipaënsis* in a 1:1 ratio (taking into account the different genome sizes). These synthetic datasets were processed identically to the progeny samples, and allele counts were calculated. Raw allele counts in the progenies were subsequently normalized using the formulas described in Lamon et al. (2025a), enabling direct comparison across samples. Normalized allele distributions for the selected BC₁ and BC₁F₂ progenies were visualized using density plots generated with the ‘ggplot2’ package in RStudio.

## RESULTS

### Development of a high-density phased peanut linkage map

A high-density phased linkage map was developed, distinguishing two phases in the BC_1_ population (Figure 3). The phase 1 map included 3,194 SNP markers, while the phase 2 map contained 6,523 SNP markers, for a total of 9,717 mapped markers (Table 1; see Supporting Information, Table S2). On average, phase 1 had 160 markers per chromosome, with the highest density observed on LG2 (234 markers). Phase 2 averaged 326 markers per chromosome, peaking at 559 markers on LG2.

**Figure 3:**
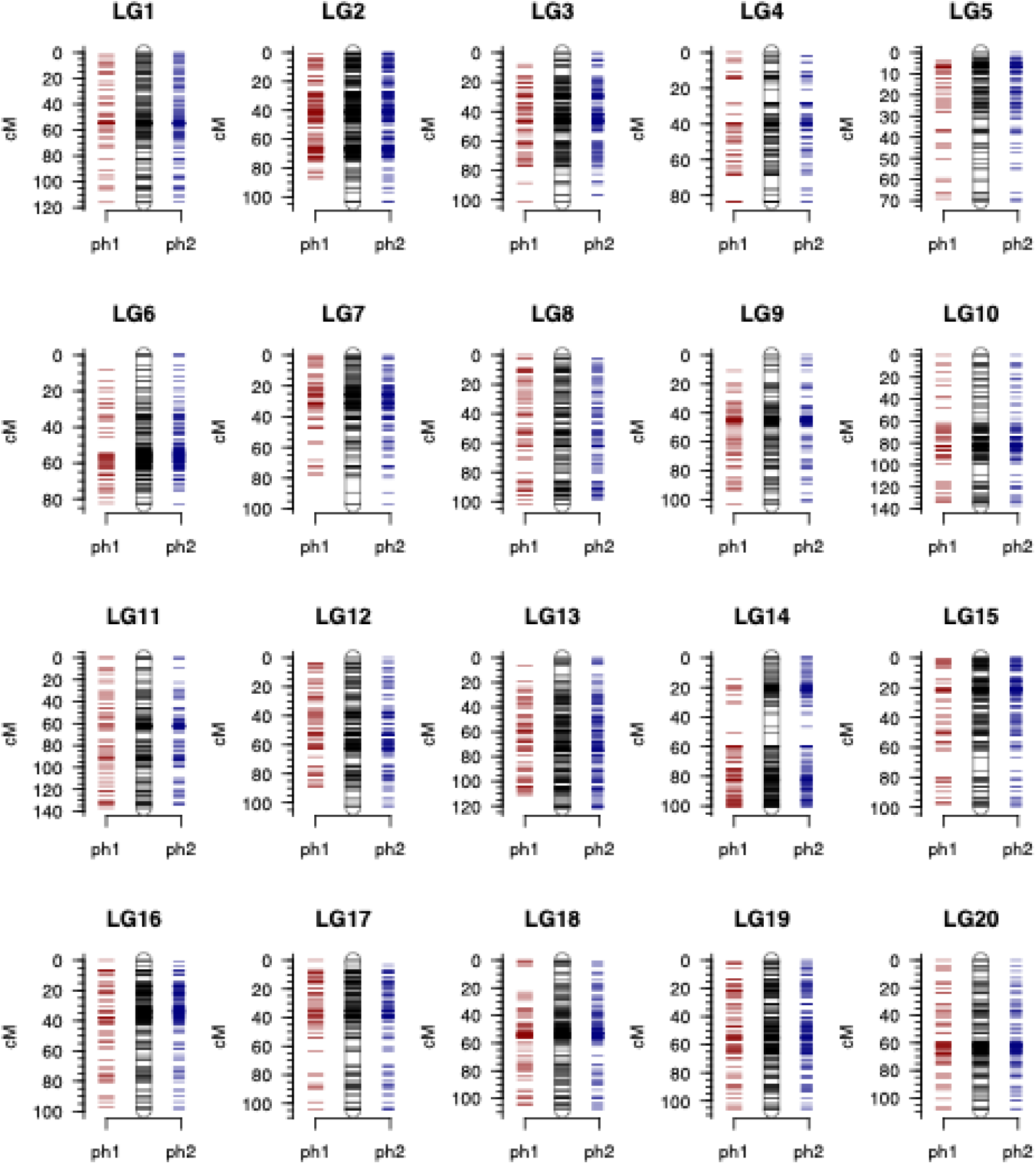
Integrated linkage map displaying 9,717 phased SNP markers (black) across 20 linkage groups (LGs 1–10 represent the A subgenome, and LGs 11–20 represent the B subgenome). Duplex × simplex markers (phase 1) are shown in red, while nulliplex × simplex markers (phase 2) are depicted in blue.

**Table 1:** Summary statistics of the integrated linkage maps, including the number of SNP markers for phase 1 and phase 2, linkage group lengths, average marker distances, and maximum marker gap distances.

| Subgenome | Linkage Group | No. of SNPs Phase 1 | No. of SNPs Phase 2 | Linkage Group Length (cM) | Average Distance (cM) | Maximum Gap Distance (cM) |
| --- | --- | --- | --- | --- | --- | --- |
| A-subgenome | LG1 | 169 | 505 | 116.11 | 0.17 | 5.65 |
|  | LG2 | 234 | 559 | 103.74 | 0.13 | 6.25 |
|  | LG3 | 174 | 345 | 101.53 | 0.2 | 6.6 |
|  | LG4 | 118 | 145 | 83.54 | 0.32 | 7.55 |
|  | LG5 | 114 | 214 | 70.24 | 0.21 | 5.08 |
|  | LG6 | 146 | 337 | 82.93 | 0.17 | 5.54 |
|  | LG7 | 182 | 343 | 97.22 | 0.19 | 9.2 |
|  | LG8 | 166 | 305 | 102.24 | 0.22 | 4.13 |
|  | LG9 | 189 | 304 | 103.71 | 0.21 | 8.52 |
|  | LG10 | 139 | 295 | 136.88 | 0.32 | 10.43 |
| B-subgenome | LG11 | 123 | 349 | 137.01 | 0.29 | 7.44 |
|  | LG12 | 127 | 244 | 103.21 | 0.28 | 5.56 |
|  | LG13 | 184 | 345 | 121.37 | 0.23 | 7.54 |
|  | LG14 | 169 | 277 | 100.94 | 0.23 | 9.04 |
|  | LG15 | 130 | 304 | 100.09 | 0.23 | 9.12 |
|  | LG16 | 157 | 298 | 99.00 | 0.22 | 5.77 |
|  | LG17 | 146 | 326 | 104.63 | 0.22 | 9.39 |
|  | LG18 | 207 | 361 | 108.44 | 0.19 | 4.41 |
|  | LG19 | 155 | 326 | 106.5 | 0.22 | 4.12 |
|  | LG20 | 165 | 341 | 108.75 | 0.21 | 4.78 |
| **Across the whole map** |  | 3194 | 6523 | 2088.08 | 0.22 | 10.43 |

The integrated map covered a total genetic distance of 2,088.08 cM, with individual LG lengths ranging from 70.24 cM (LG5) to 137.01 cM (LG11). The average marker interval was 0.22 cM, while maximum distances between markers varied from 4.12 cM (LG19) to 10.43 cM (LG10). Strong cross-LG signals were found between LG3 and LG13, LG4 and LG14, and LG9 and LG19 (Figure S6).

Marey plots displayed distinct S-shapes with smooth transitions (Figure 4A), indicative of a high-quality linkage map and consistent with expected recombination rate variation along the chromosomes. An exception was observed for LG5, which showed near-complete suppression of recombination along its upper arm. This pattern is consistent with the presence of a large post-polyploidy inversion in this region of *A. hypogaea* (Bertioli et al. 2019), an inversion that likely stabilized the homeologous chromosome pair by disrupting collinearity between the A and B subgenomes and would also be expected to suppress recombination between cultivated and wild chromosome 05 homologs. In addition, LG4 and LG14 showed extreme “doubled-line” artifacts.

**Figure 4:**
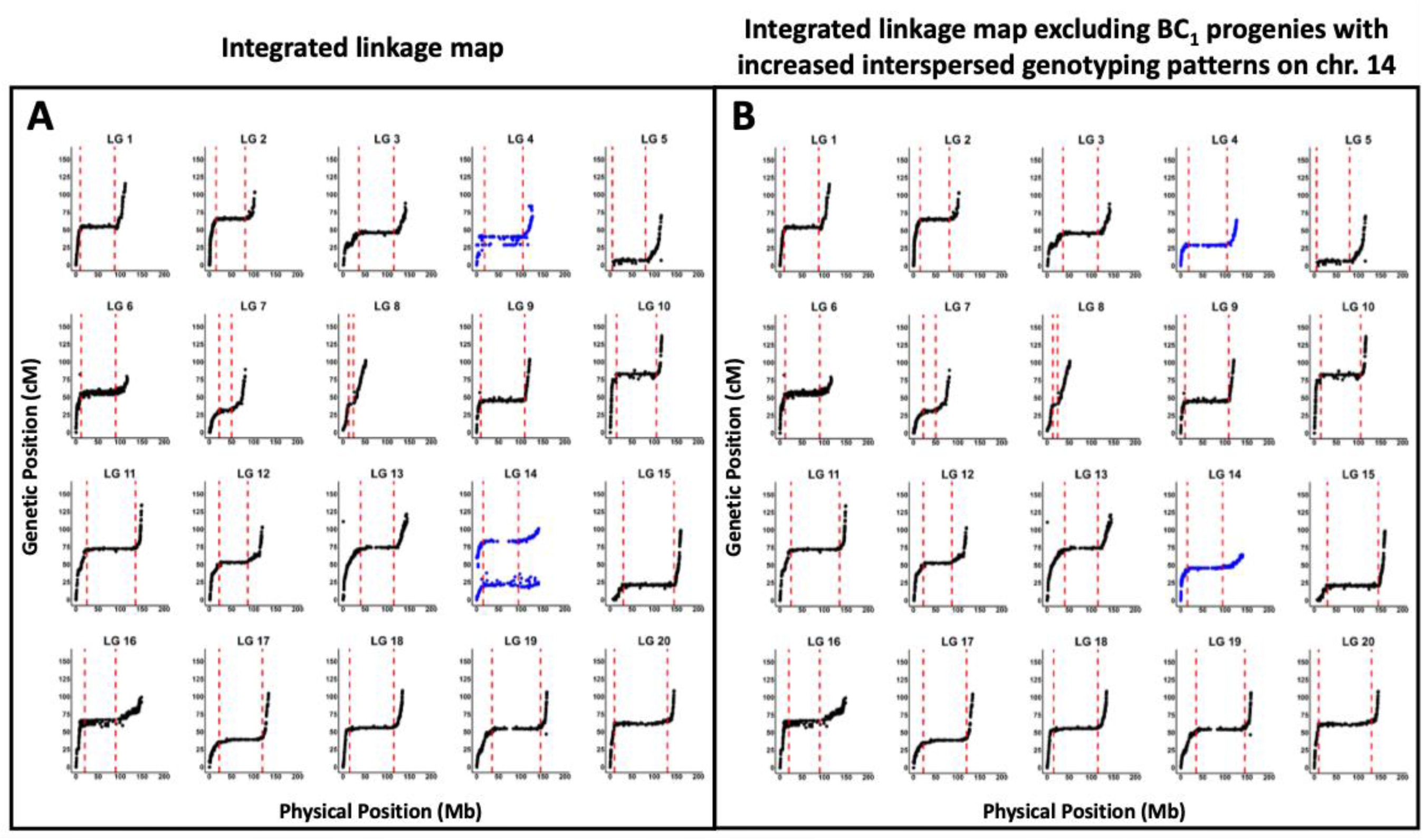
Marey plots of the integrated linkage map (A) and the map after excluding BC_1_ progenies with increased interspersed genotyping patterns on linkage group (LG) 14 (B) are shown. The y-axis represents genetic positions, and the x-axis corresponds to physical positions in the ‘Tifrunner’ reference genome. In panel A, double LG artifacts are evident, particularly on LGs 4 and 14 (blue). These artifacts disappear in panel B following the exclusion of the affected progenies. Pericentromeric regions are indicated by red dashed lines.

LG14 was particularly affected, presenting as two complete S-shaped curves.

### Genotyping and sequencing analyses reveal a correlation between interspersed genotyping patterns and unbalanced genomic compositions, which significantly affect linkage mapping

In this section, we describe how we discovered that two initially unexpected observations—chromosome-scale interspersed SNP genotyping patterns and the appearance of “doubled” LGs—were related, and we identify their underlying cause.

Regions with interspersed genotyping patterns were most frequently observed in chromosome set (LGs 4 and 14), followed by sets 2 (LGs 2 and 12), 6 (LGs 6 and 16), 3 (LGs 3 and 13), 5 (LGs and 15), and 9 (LGs 9 and 19), in descending order of frequency (Figure S3B, Figure S7, A and B). Overall, phase 1 exhibited substantially fewer interspersed genotyping patterns than phase 2, although it still showed a notable level of background noise. Notably, the LGs containing the highest numbers of individuals with interspersed genotyping patterns also showed the most extreme “doubled” LG behavior.

With this in mind, we followed the intuition that these chromosome-scale interspersed genotyping patterns represented some chromosomal entity, or biological circumstance, that did not follow standard assumptions, and that this might underlie the doubled LG patterns. To test this, 18 BC_1_ progenies with pronounced interspersed genotyping on LG14 in the phase 2 map were excluded, and linkage mapping was repeated. Satisfyingly, this eliminated the doubled LG patterns (Figure 4B, Figure S3A).

Then, to investigate the nature of the entity or circumstance underlying the interspersed genotyping patterns, we performed WGS on BC_1_ progenies with (Ind 47 and 62) and without (Ind 33 and 39) the interspersed genotyping patterns in LG14. In addition, BC_1_F_2_ progenies Ind 40.3 and 40.4, descendants of BC_1_ individual Ind 40, which also exhibited interspersed genotyping on LG14, were sequenced.

Sequencing revealed that individuals with interspersed genotyping patterns had unbalanced genomic compositions on the same chromosomes. Specifically, AAAB configurations were detected in chromosome set 4 in Ind 47 and 62 that displayed interspersed genotyping, but not in Ind 33 and 39 that had normal genotyping patters (Figure 5, A–D). Similarly, BC_1_F_2_ progenies Ind 40.3 and 40.4 showed the same large unbalanced AAAB configuration in chromosome set 4 (Figure 5, E and F).

**Figure 5:**
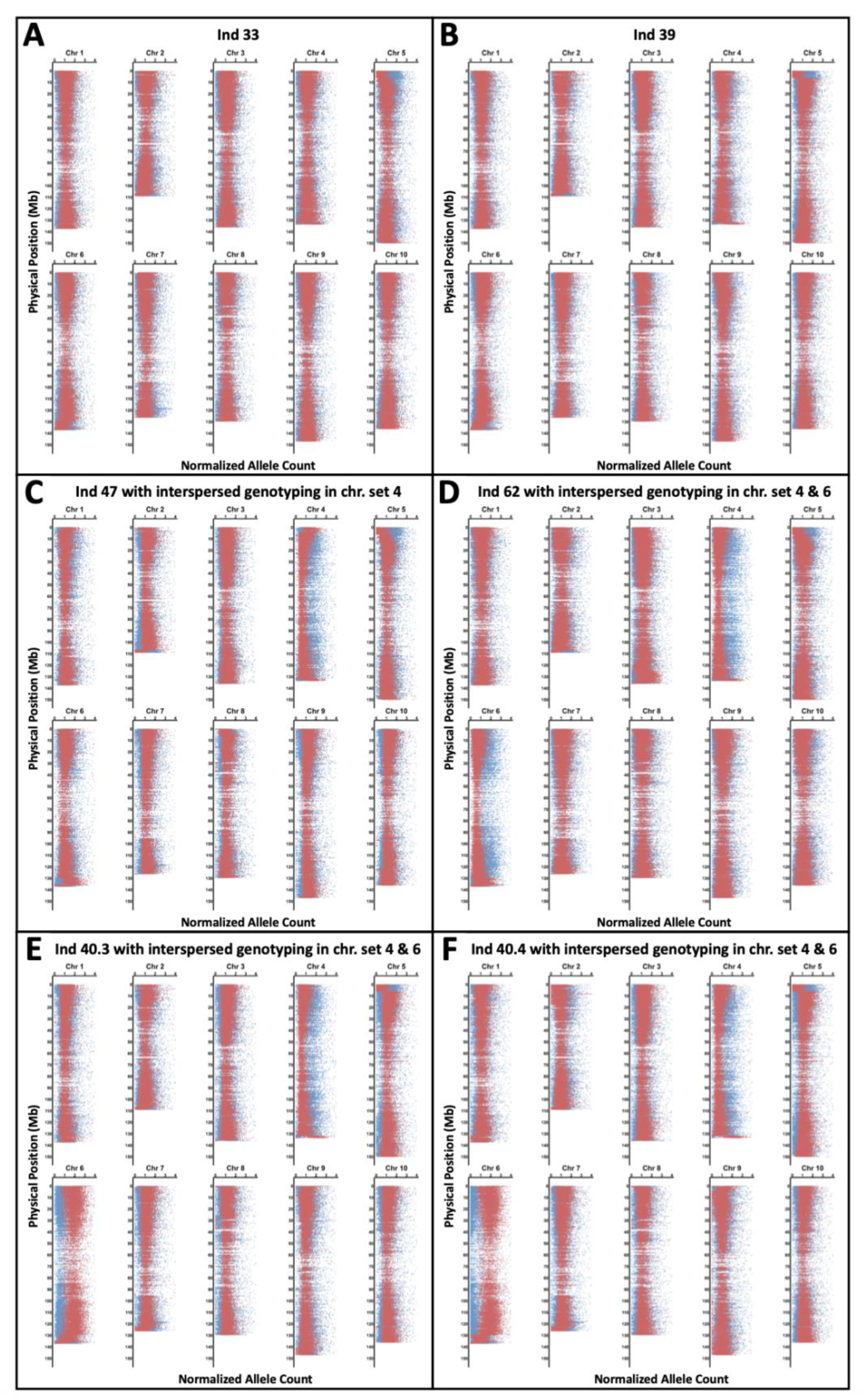
Density plots illustrating read counts by chromosome for sequenced genotypes distinguish between *Arachis duranensis*-like (AA-genome; blue) and *A. ipaënsis*-like (BB-genome; red) subgenomes. Chromosomes with a balanced composition of A and B subgenomes (AABB genome) show read counts of 1 for both A and B reads. In contrast, chromosomes with genome shifts towards either *A. duranensis* or *A. ipaënsis* display varying values, with common scenarios being A = 1.5 and B = 0.5 for the AAAB-genome, and A = 0.5 and B = 1.5 for the ABBB-genome. Panels A and B show Ind 33 and Ind 39, both with regular (balanced) chromosome set 4. Panel C displays Ind 47, which has an unbalanced chromosome set 4, consistent with the presence of interspersed genotyping patterns. Panel D shows Ind 62, which exhibits unbalanced chromosome sets 4 and 6, also reflecting the interspersed genotyping patterns observed in genotyping analysis. Panels E and F present BC_1_F_2_ progenies Ind 40.3 and 40.4, both derived from Ind 40, each displaying unbalanced chromosome sets 4 and 6 as predicted from genotyping analysis.

Across all individuals, chromosome sets 2–7 exhibited individual-specific genomic composition variation, whereas sets 1, 8, 9, and 10 were largely balanced. Notably, individuals with an AAAB composition at the top of chromosome set 5 had yellow flowers (Figure S8). This is in agreement with a previous report of the genetic loci determining flower color, (Bertioli et al. 2019), and the dominance of the yellow wild allele carried on the B subgenome in this population.

### DR estimation in segmental allotetraploid peanut

Pericentromeric boundaries for each LG were visually estimated to allow clearer interpretation of DR event distribution (Figure 4, A and B; see Supporting Information, Table S3). In both phase 1 and phase 2, DR scores were predominantly detected near the telomeric regions, consistent with previous studies (see Supporting Information, Table S4) (Bourke et al. 2015). Specifically, in phase 1, DR events occurred at the top of LG7 (0.34–6.54 Mb, Ind 21), the top of LG12 (1.49–1.77 Mb, Ind 38), the bottom of LG15 (157.87–159.92 Mb, Ind 59), and the top of LG19 (0.18–0.43 Mb, Ind 59), with all physical positions referenced to the ‘Tifrunner’ genome (Figure 6A). In phase 2, DR events were consistently observed at the top of LG4 (1.94–3.88 Mb in the ‘Tifrunner’ genome) in six individuals (Ind 8, 11, 40, 47, 58, and 59) (Figure 6B). Thus, DR events were most frequent on LG4 in phase 2, whereas in phase 1 they were distributed across several LGs but always confined to telomeric regions.

**Figure 6:**
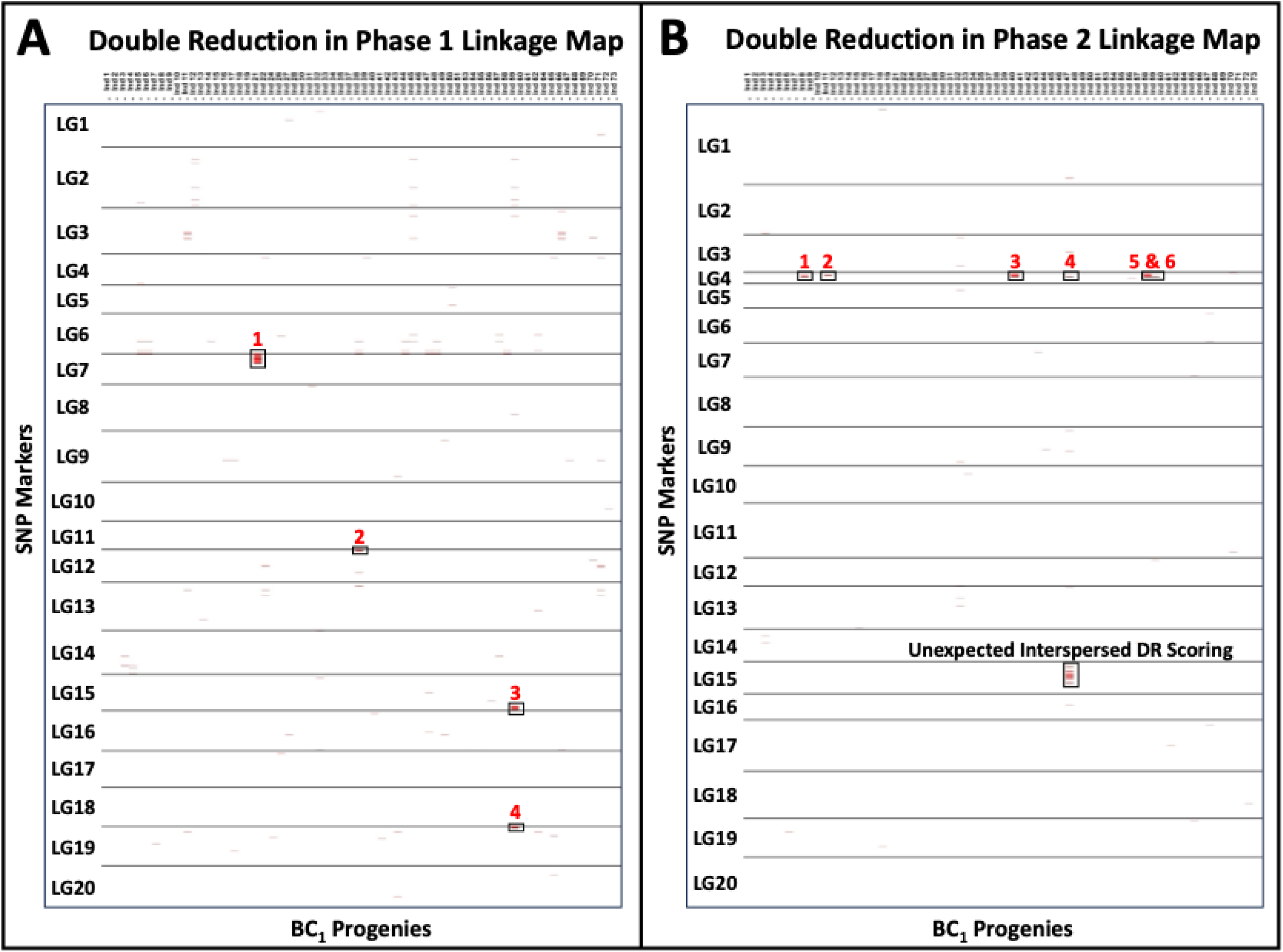
Heatmaps display double reduction (DR) events in BC_1_ progenies based on phase 1 (A) and phase 2 (B) linkage maps, with SNP markers ordered by genetic distance. Red marks indicate SNP scores consistent with DR, while regions with consecutive DR-consistent scores, identified as DR events, are highlighted with black rectangles and numbered in red. In agreement with previous reports, DR events occurred either at the top or bottom of the linkage groups: the top of LG7 (1), top of LG12 (2), bottom of LG15 (3), and top of LG19 (4) in the phase 1 map, and the top of LG4 (1–6) in the phase 2 map. In phase 2, unexpected interspersed DR scoring was observed across LG15 in Ind 47.

Among the BC_1_ progenies, 12% (8 out of 66) showed evidence of DR in either phase 1 or phase 2 linkage maps (see Supporting Information, Table S4). Because only a subset of marker combinations was analyzed, this figure likely underestimates the true frequency of DR. In addition, an AAAB genomic configuration was detected in the sequenced individual Ind 47 within the DR region (Figure 5C, Figure 7). This result confirms the theoretical expectation that DR events generate unbalanced genomic composition in progenies (Figure 1).

**Figure 7:**
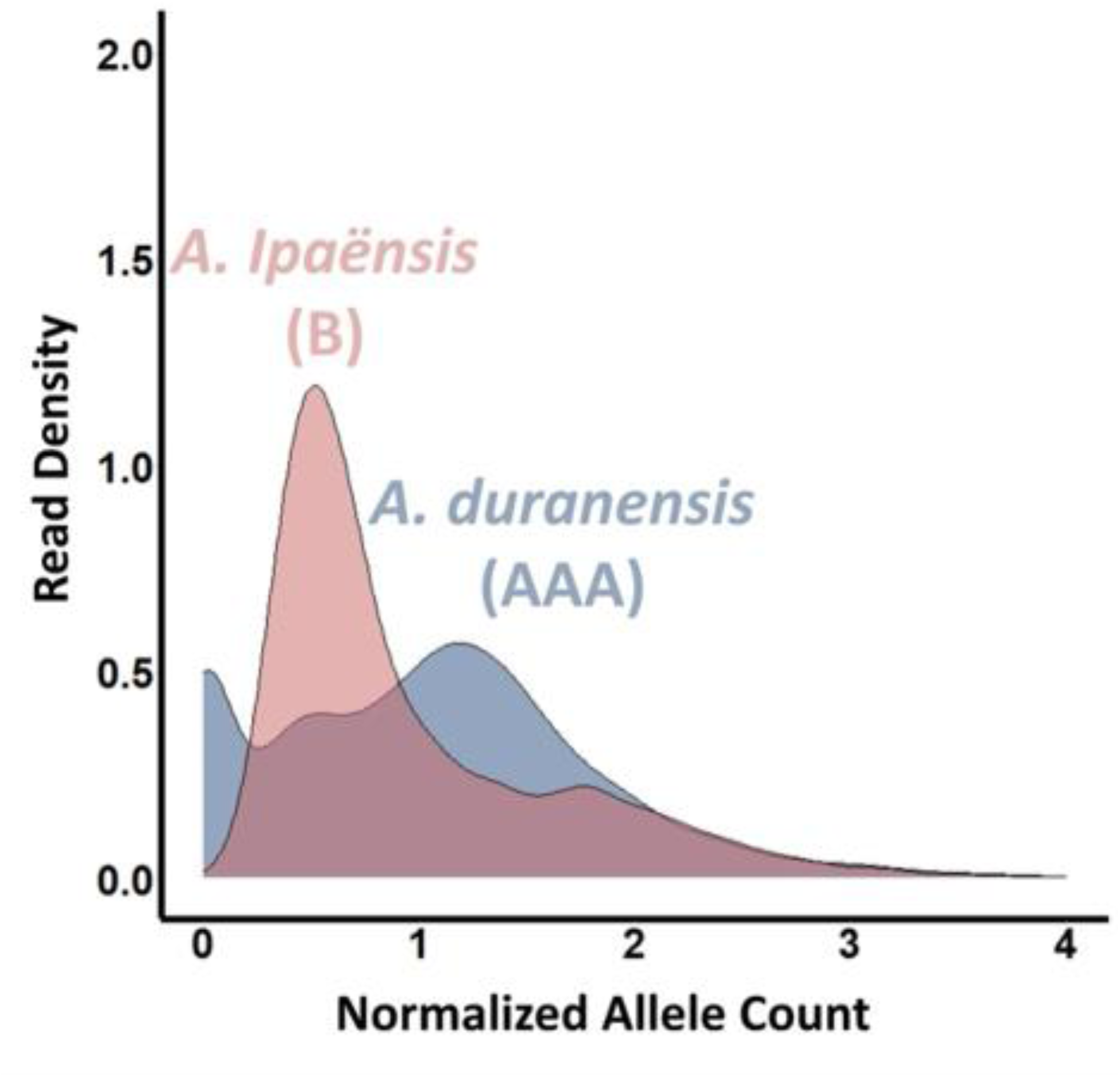
Normalized allele count read density plot for the upper region of LG4 (3.15–3.88 Mb in the ‘Tifrunner’ genome; 4.27–4.97 Mb in *Arachis ipaënsis* gnm1) of individual 47, where double reduction duplex scores were detected. The expected genome composition for this region is T^a^T^a^G^a^G^b^, where “T” and “G” are arbitrary alleles, with the T^a^ allele provided by *A. stenosperma*, the G^a^ allele from the A subgenome of ‘IAC OL4’, and the G^b^ allele from the B subgenome of ‘IAC OL4’. The plot demonstrates a shift in genome composition toward *A. duranensis* (AAAB), as indicated by a normalized allele count of approximately 1.5 for the A subgenome and 0.5 for the B subgenome. This scenario corresponds to the genomic configuration depicted in Figure 2, C and D.

However, Ind 47 also displayed an unexpected interspersed pattern of DR scores across LG15, including substantial presence in the pericentromeric region, an area typically not associated with DR (Bourke et al. 2015). Notably, Ind 47 did not exhibit the anticipated unbalanced genomic configuration in this region (Figure 5C), suggesting that a different genetic mechanism, rather than DR, may underlie these patterns.

### Hidden Markov model revealed homoeologous exchange but could not confirm DR

The Hidden Markov Model using a tetrasomic model allowed for both bivalent and multivalent pairing. The model was run at the default error prior setting (Figure S9, Figure S10). With these setting, two individuals were predicted to have arisen from multivalent pairing in the parental meioses, namely Ind 47 on chromosome 05 and Ind 21 on chromosome 07 (Figure S11). However, when the process was repeated over a range of possible error priors, the maximum likelihood results no longer predicted multivalent pairing for these (or any other) individuals (Figure S11). However, there were six individuals that were predicted to have arisen from homoeologous exchanges on chromosomes 2, 3, 5 and 7 (Figure S12).

## Discussion

In this study, we estimated DR occurred in 12% of the analyzed BC_1_ peanut progenies, a frequency that is unexpectedly high for a segmental allotetraploid species. This observation is consistent with previous findings in peanut (Leal-Bertioli et al. 2015a), which reported elevated levels of tetrasomic inheritance, despite the predominance of disomic inheritance. While DR frequencies in autopolyploids are generally higher due to their higher rate of multivalent formation, our results suggest that, in peanut, DR is still a rare but recurrent event and can be reliably estimated when sufficient genomic resolution and appropriate population structures are used.

However, it is important to recognize that several methodological limitations likely resulted in an underestimation of DR rates in our study. For example, the number of informative marker combinations available for estimating DR was limited, and stringent filtering of marker dosages was necessary to ensure accurate segregation scoring. Additionally, the inherent structure of the BC_1_ population only permits estimation of certain DR events. Alternative mechanisms, such as tetrasomic inheritance in the female parent in phase 1 or consecutive rounds of tetrasomic inheritance in the male parent in phase 2, could also generate genotypic patterns that resemble DR, potentially leading to overestimation. Nevertheless, our conservative scoring approach, which required multiple consecutive markers with consistent dosage, minimizes the risk of false positives and strengthens confidence in the DR events we report.

Most notably, our results provide direct evidence that DR, which depends on homoeologous exchange, could lead to an increase of unbalanced genomic compositions, such as ABBB and AAAB states, in the progeny of allopolyploid species like peanut (Figure 1, Figure 7). These findings establish a new theoretical framework for estimating DR in allopolyploids by demonstrating that the presence of ABBB and AAAB genome compositions can serve as a possible indicator of DR events.

All identified DR events in our study were localized to telomeric regions, in agreement with established theory and previous empirical observations (Mather 1936; Fisher 1947; Haynes and Douches 1993; Butruille and Boiteux 2000; Stift et al. 2008; Nemorin et al. 2012; Bourke et al. 2015). This localization likely reflects higher crossover frequency and multivalent formation near telomeres compared to the more structurally constrained pericentromeric regions. Interestingly, we observed an unusual interspersed pattern of DR-like scores across the pericentromeric region of LG15 in Ind 47, which was not associated with an unbalanced genomic configuration in the corresponding chromosome set. This finding suggests other genetic phenomena could underlie these observations. Ind 47 was also initially predicted to have arisen from DR using the HMM, although this prediction was subsequently lost at a higher error prior. However, the autotetraploid HMM we employed is reliant on a high-quality integrated linkage map. Our pseudo-integrated map was visibly discordant on certain LGs including chromosome 05 (Figure S5), which may have affected the quality of the results. Therefore, determining presence or absence of unbalanced genomic compositions could become useful for distinguishing true DR events from other genotypic patterns in allopolyploid mapping populations.

The generation and analysis of a BC_1_ population was critical for uncovering these findings. Because peanut is highly self-pollinating, establishing a backcross population requires substantial manual effort. However, this approach was essential for generating the segregation patterns needed to detect the transition from a single dosage to duplex gametes (Moretzsohn et al. 2023). Furthermore, the development of the BC_1_ population allowed us to construct a high-density phased linkage map for peanut, using SNP chip technology and the ‘fitPoly’ and ‘polymapR’ packages in RStudio.

This map achieved high marker density compared to previous studies and proved crucial for estimating DR (Zhou et al. 2014; Leal-Bertioli et al. 2015b; Huang et al. 2016; Li et al. 2017; Han et al. 2018; Ballén-Taborda et al. 2019; Liu et al. 2020; de Blas et al. 2021; Sun et al. 2022; Miao et al. 2023; Tsai et al. 2024). High marker density enhances both the sensitivity and accuracy of DR estimation, as well as the ability to resolve complex patterns of recombination and genome composition in allopolyploid backgrounds.

Our mapping and sequencing analyses further revealed that regions with frequent interspersed genotyping patterns, especially in LG14, coincided with unbalanced genomic compositions and double LGs artifacts in Marey plots. In some cases, these mapping artifacts were so extensive that they split a single LG into two distinct segments. Although similar issues have been reported informally in previous studies, they have rarely been addressed in detail in the literature. Commonly, researchers have focused on the primary LG and excluded alternative signals to simplify map interpretation. While this practice improves clarity, it also reduces genetic resolution and may overlook important aspects of genetic instability.

The distortions caused by unbalanced genome composition and DR are likely to remain a recurring challenge in genetic mapping of peanut and other allopolyploid organisms. Unbalanced configurations resulting from homoeologous exchange or DR not only complicate map construction, but also affect the interpretation of genetic diversity and inheritance patterns in breeding populations (Lamon et al. 2025b). This highlights the need for analytical tools and pipelines that can detect and account for these genomic features during linkage analysis.

To address these challenges, future mapping strategies in allopolyploids should incorporate genome composition data, ideally through sequencing or improved dosage scoring, as early as possible in map construction. As demonstrated in this study, the systematic identification and filtering of hotspots for interspersed genotyping prior to map building, combined with sequencing to confirm the presence and boundaries of unbalanced genomic segments, can help prevent mapping artifacts and improve the accuracy of both map construction and downstream analyses.

Another option is to improve genotyping software so that it can dynamically accommodate the complexity of allopolyploid genome structures. For example, in peanut, in genomic regions where frequent multivalent formation and recombination between divergent subgenomes are expected, standard tetraploid scoring (0–4 scale) may not be sufficient to represent all possible allele dosages. Software that allows a more granular and context-dependent dosage scale, coupled with linkage mapping tools capable of handling such data, would be particularly valuable for peanut and other segmental allopolyploids.

Beyond their technical implications, our findings have evolutionary significance. The estimation of DR as a rare but consistent form of genetic instability in cultivated peanut provides direct evidence for the ongoing generation of new allelic combinations, even in crops with narrow genetic bottlenecks and a high degree of subgenome fixation. Both DR and homoeologous exchange contribute to genome plasticity, offering a partial escape from the genetic constraints imposed by polyploidization bottleneck and domestication. These mechanisms may have played an important role in the morphological diversification and adaptability that have characterized peanut’s evolutionary history.

In summary, this study estimated DR, a rare but biologically meaningful outcome of genetic instability in peanut, which depends on homoeologous exchange and multivalent formation. Our findings underscore the value of high-density linkage mapping in detecting and interpreting rare meiotic events and call for the continued development of analytical approaches that can accurately accommodate the genomic complexity of segmental allopolyploids. Addressing these challenges will be essential for improving the precision of genetic studies and breeding strategies not only in peanut, but also in other important allopolyploid crops.

## Conclusions

This study presents a high-density phased linkage map for peanut, constructed using 9,717 SNP markers. Unbalanced genomic compositions were identified in several BC_1_ progenies, mainly caused by homoeologous exchange, which were shown to reduce map quality and introduce artifacts such as double LGs. Excluding progeny with extensive genomic imbalance improved overall map resolution and accuracy. Evidence of DR was found in 12% of the BC_1_ population progenies, across LGs 4, 7, 12, 15, and 19. Of particular interest, a DR event on LG4 corresponded with an AAAB genome configuration, lending empirical support to the hypothesis that DR can result in unbalanced chromosome sets in segmental allopolyploids like peanut. The ability to estimate such rare events has significant implications for understanding the evolution and genetic complexity of allopolyploid organisms.

## Data availability statement

WGS data are currently under embargo and will be released upon publication. Sequencing analysis scripts are available at https://github.com/brianabernathy/ABGD. All other data generated or analyzed during this study are included in the supporting information.

## Acknowledgments

Special thanks are extended to Adriana Regina Custodio, Jennifer Leverett, and Mark Hopkins for their valuable contributions. Samuele Lamon also acknowledges the Institute of Plant Breeding, Genetics, and Genomics at the University of Georgia for the 2023 Peggy Ozias-Akins Leadership Award.

## Author Contributions

Samuele Lamon conducted the genotypic data analysis, including linkage mapping, identification of interspersed genotyping patterns, and estimation of double reduction. He also performed sequencing data visualization and co-wrote the manuscript. Peter Bourke assisted with genotypic data analysis, linkage mapping, and estimation of double reduction, and contributed to manuscript preparation. Brian Abernathy oversaw the sequencing data analysis. João Francisco dos Santos and Ignácio José de Godoy developed and provided the plant material for the study. Soraya Leal-Bertioli conceptualized the project. David Bertioli conceptualized the project, contributed to sequencing data analysis, and co-wrote the manuscript.

---

**Figure S1:**
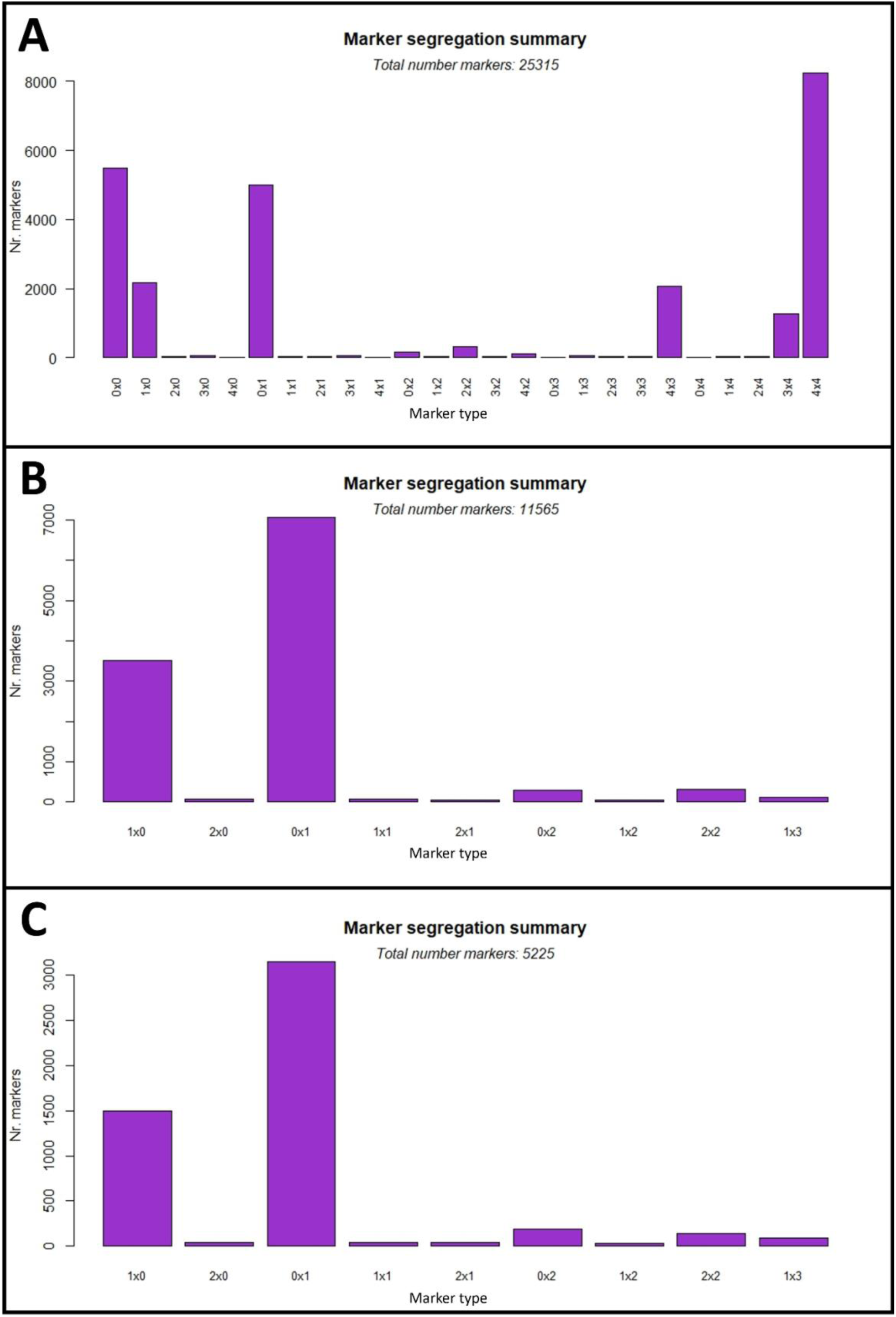
Panel A shows the segregation of markers after filtering for non-missing values in both ‘IAC OL4’ and MagSten, retaining 25,315 markers from the BC_1_ population, with MagSten as the donor and ‘IAC OL4’ as the recurrent parent. Panel B displays the segregation after converting marker combinations to basic types and removing non-segregating markers, resulting in 11,565 retained markers. In panel C, the segregation is shown after removing non-segregating markers and screening for duplicates, yielding 5,225 retained markers. The x-axis represents marker combinations: nulliplex (0), simplex (1), duplex (2), triplex (3), and tetraplex (4).

**Figure S2:**
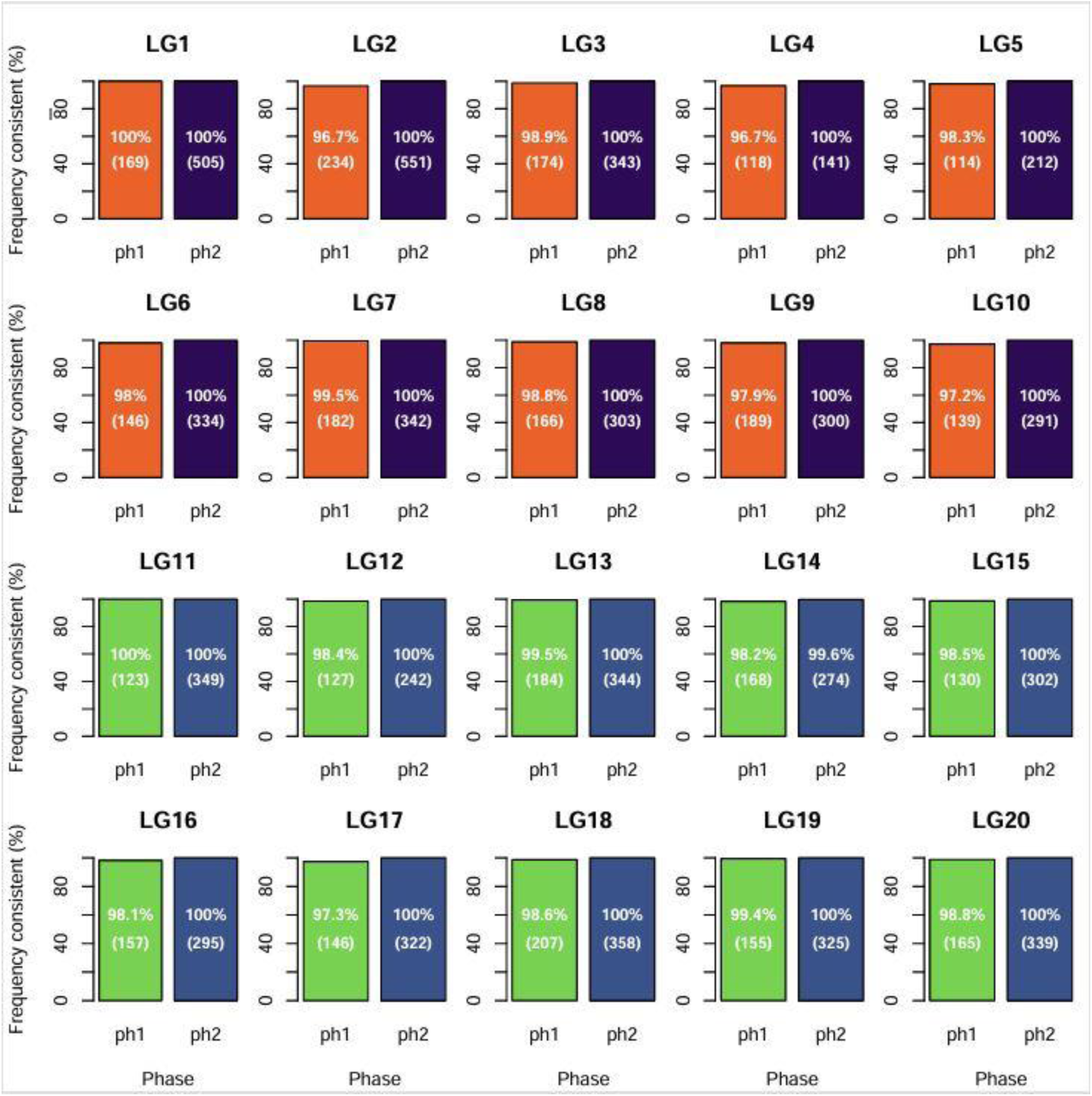
Bar plots of phasing consistency by linkage group (LG), where LGs 1–10 correspond to the A subgenome and LGs 11–20 correspond to the B subgenome. Phase 1 is shown in orange (A subgenome) or green (B subgenome), and phase 2 is shown in dark blue (A subgenome) or light blue (B subgenome). The y-axis indicates the percentage of markers whose phasing from genotype calling matches the linkage mapping phasing, with the total number of markers given in parentheses.

**Figure S3:**
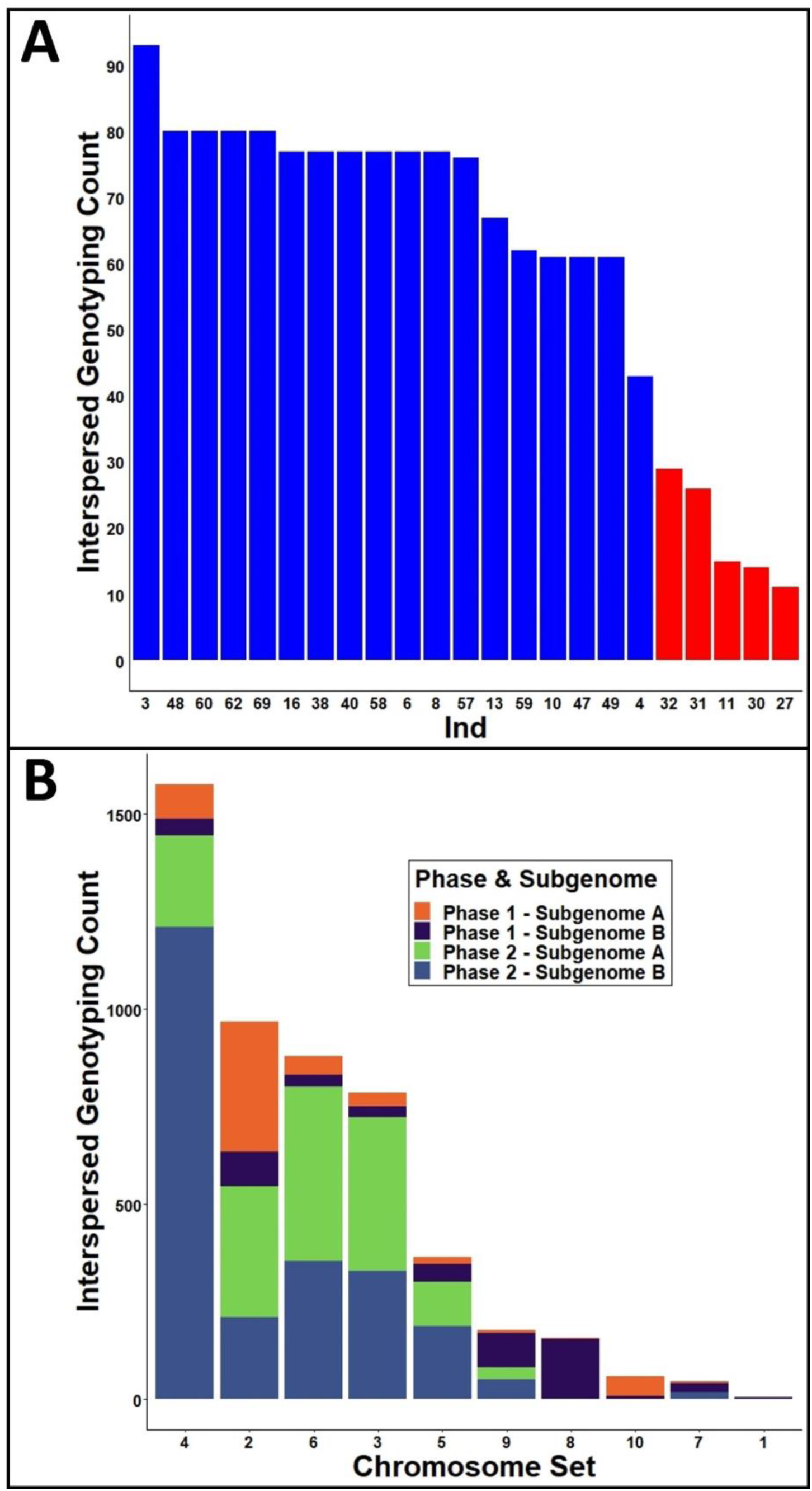
Distribution and genomic context of interspersed genotyping events detected in BC_1_ progenies. In panel A, bar plot displaying 18 BC_1_ individuals (Ind) with interspersed genotyping events specifically on LG14 (corresponding to chromosome set 4 subgenome B) of phase 2. Individuals with increased counts are shown in blue, while those with reduced counts are shown in red. Progenies with no interspersed events are excluded. In panel B, stacked bar plot summarizing total interspersed genotyping counts across all chromosome sets. Each bar is subdivided by phase (1 or 2) and subgenome (A or B), as indicated in the legend.

**Figure S4:**
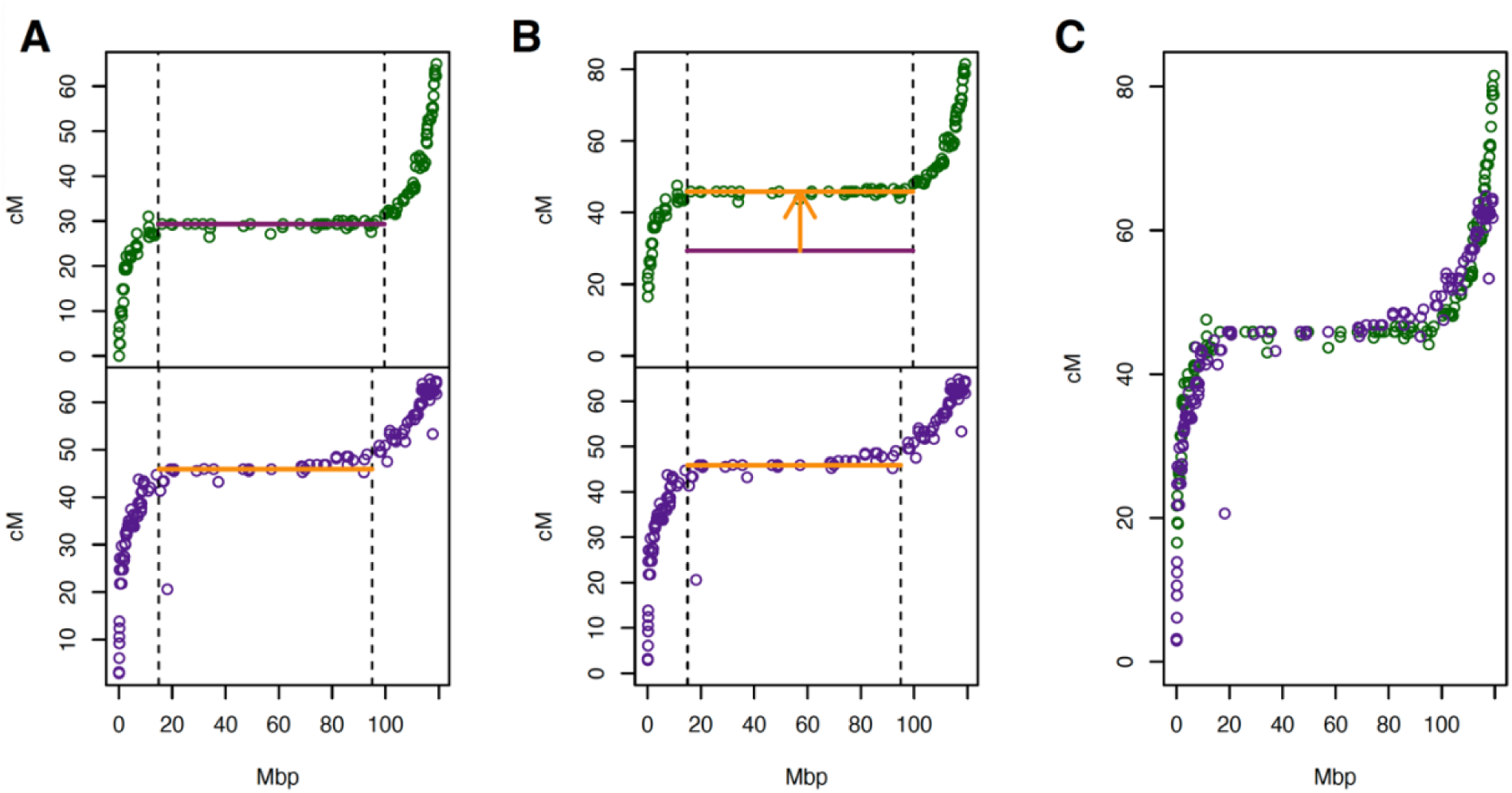
Steps in the generation of pseudo-integrated linkage map of chromosome 04. In panel A, centromeric bounds were visually identified (vertical dashed lines) and the genetic position of the centromeres were estimated as the median cM position of markers within this interval. Upper plot shows linkage group (LG) 4 (green points) while the lower plot shows LG14 (purple points). In panel B, markers on LG4 were shifted upwards by 16.5 cM, the distance between the two centromeric lines (denoted by arrow). Panel C shows the pseudo-integrated map with centromeres aligned (green points for LG4 markers and purple for LG14).

**Figure S5:**
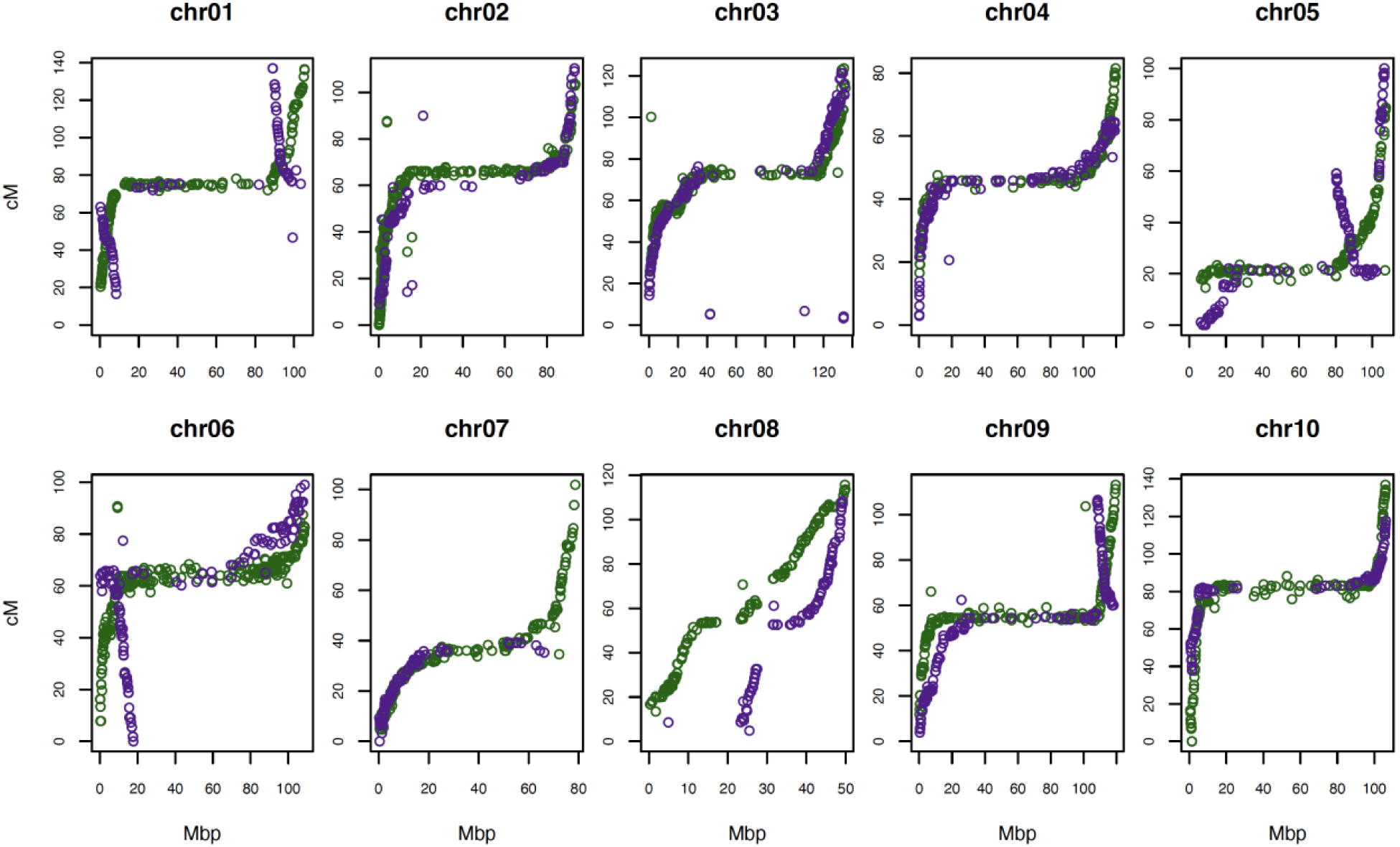
Pseudo-integrated linkage maps. Linkage group (LG) 1 – LG10 are shown in green, while LG11 – LG20 are shown in purple.

**Figure S6:**
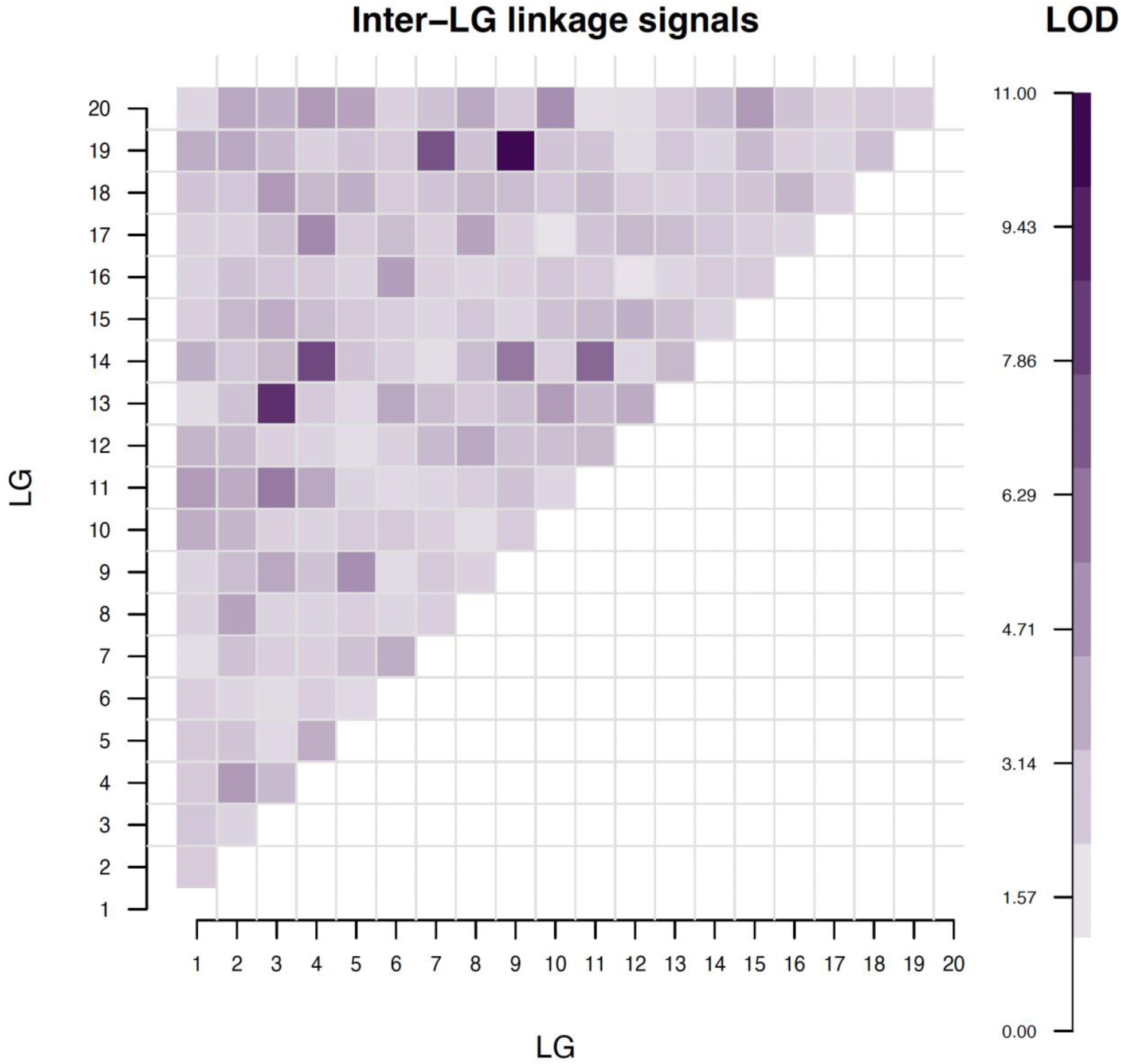
Heatmap of the maximum LOD for linkage detected between pairs of markers located on different linkage groups (LGs). The strongest cross-LG signals were found between LG3 – LG13 (chromosome 03), LG4 – LG14 (chromosome 04), and LG9 – LG19 (chromosome 09) shown in darker shades of purple. The pairs of markers involved in generating these maximum LOD scores were “well-linked” within their respective LGs, suggesting that the linkage signals were not due to mis-assigned markers. In these three groups, multiple other cross-LG linkages at elevated LOD scores were also detected.

**Figure S7:**
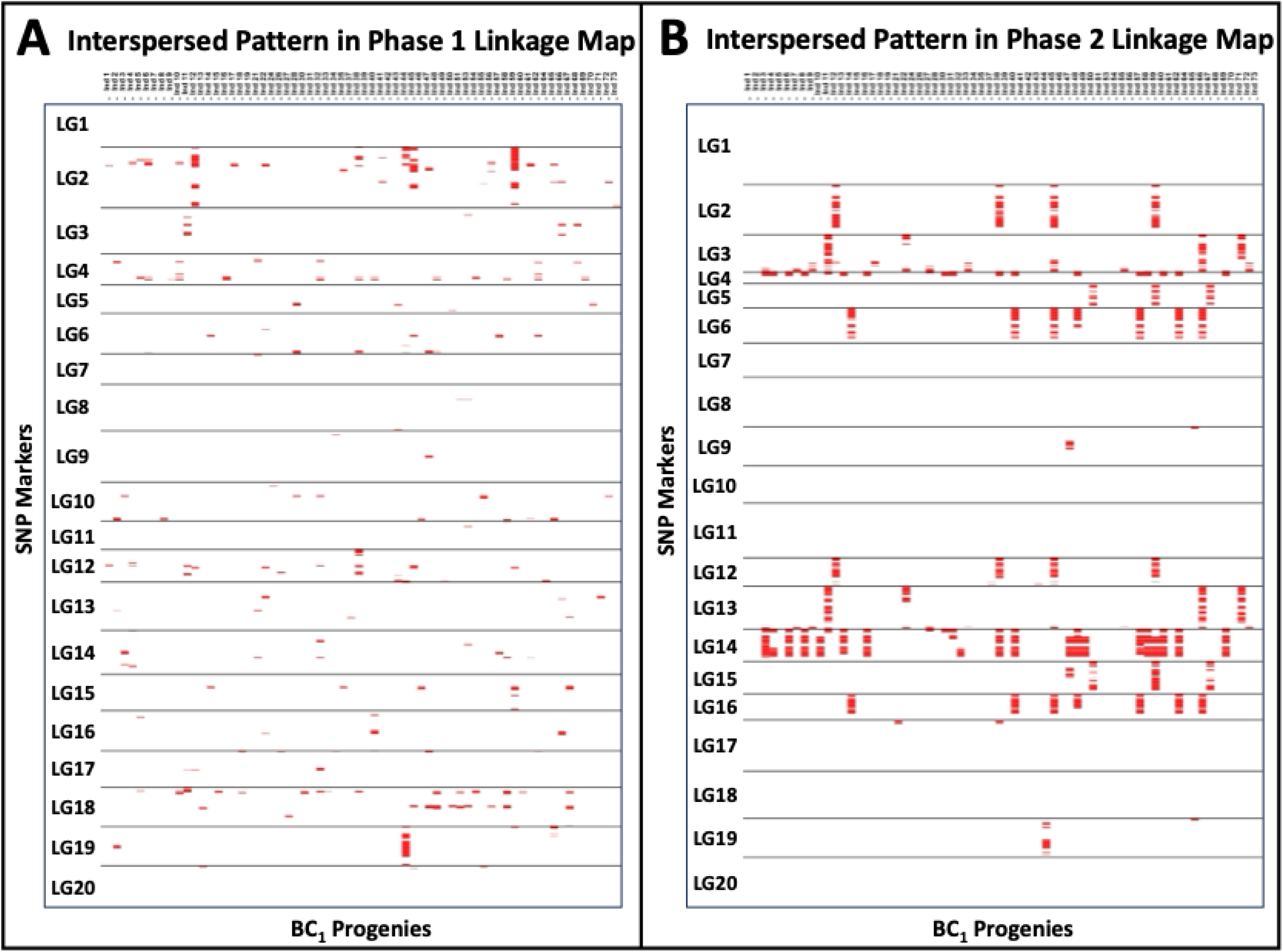
Heatmaps displaying interspersed genotyping patterns (in red) in BC_1_ progenies based on phase 1 (A) and phase 2 (B) linkage maps. SNP markers are ordered by ‘Tifrunner’ physical distance. In panel A, phase 1 shows generally low levels of interspersed patterns, with scattered noise and localized concentrations in linkage group (LG) 2, LG12, and LG19. In panel B, phase 2 exhibits a markedly higher prevalence of interspersed genotyping, with pronounced patterns observed across homologous chromosome sets: LG2 and LG12, LG3 and LG13, LG4 and LG14, LG5 and LG15, LG6 and LG16, and LG9 and LG19.

**Figure S8:**
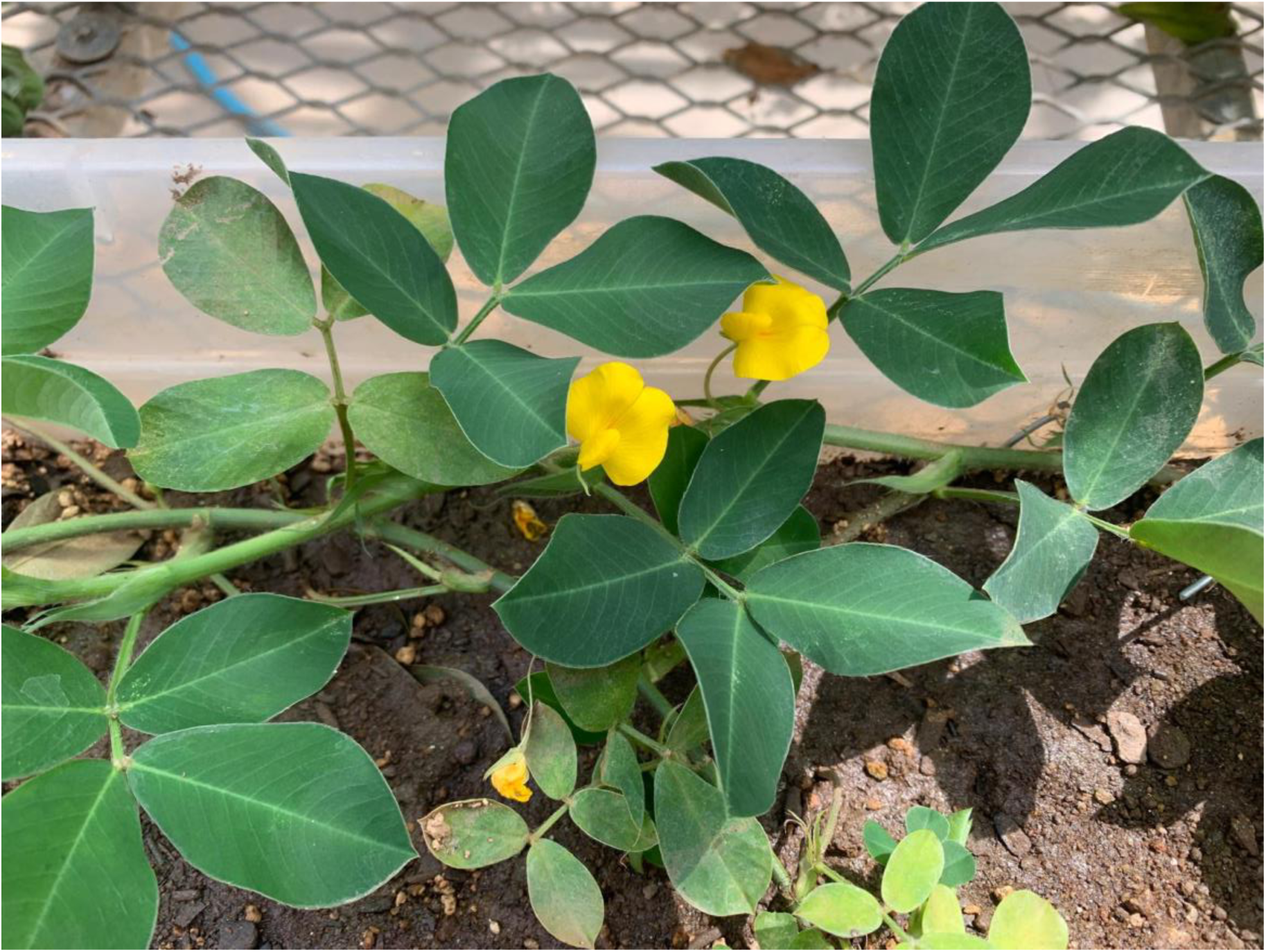
BC_1_ and BC_1_F_2_ derived from a cross between the neoallotetraploid [*Arachis magna* K 30097 x *A. stenosperma* V 15076]4x (MagSten) and ‘IAC OL4’ exhibited yellow flower phenotypes.

**Figure S9:**
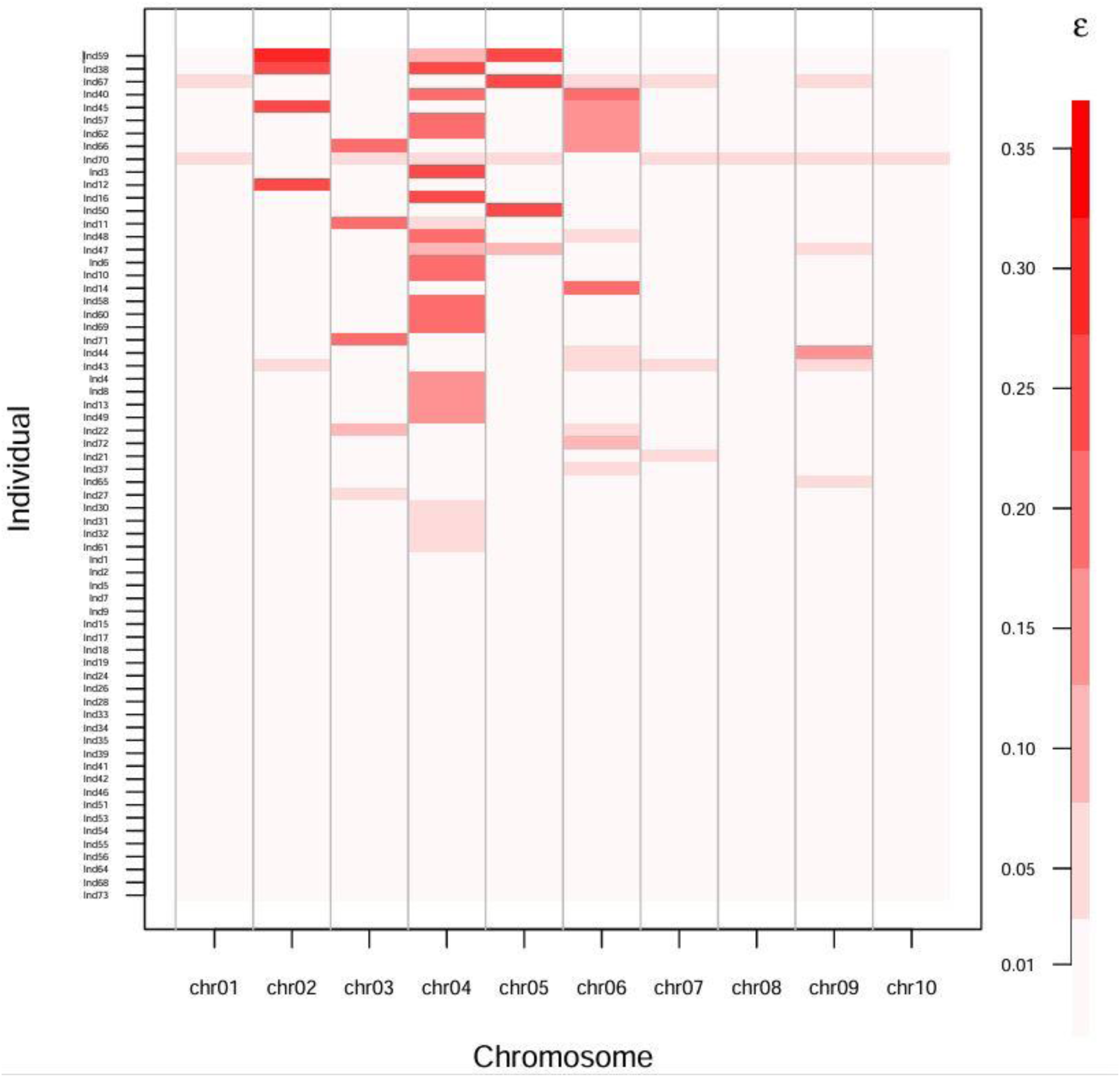
Heatmap showing the predicted genotyping error rates for each individual (y-axis) and chromosome (x-axis) combination, using a tetraploid HMM run over a range of error priors (ranging from 0.01 – 0.35 as shown in the color-bar on the right).

**Figure S10:**
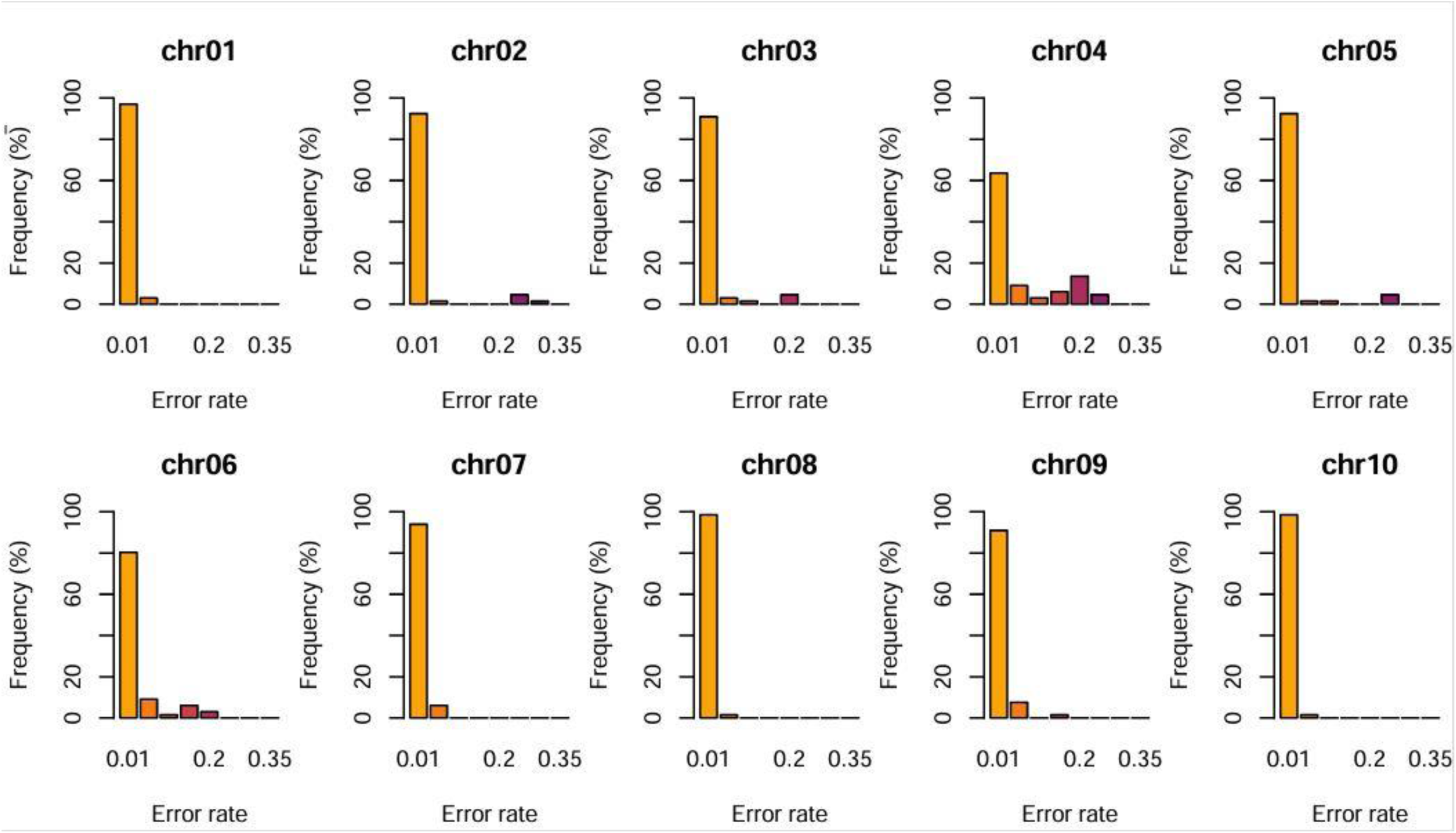
Distribution (histogram) of the predicted genotyping error rates per chromosome using a tetraploid HMM run over a range of error priors (ranging from 0.01 – 0.35). The underlying data is the same as that used to generate Figure S9.

**Figure S11:**
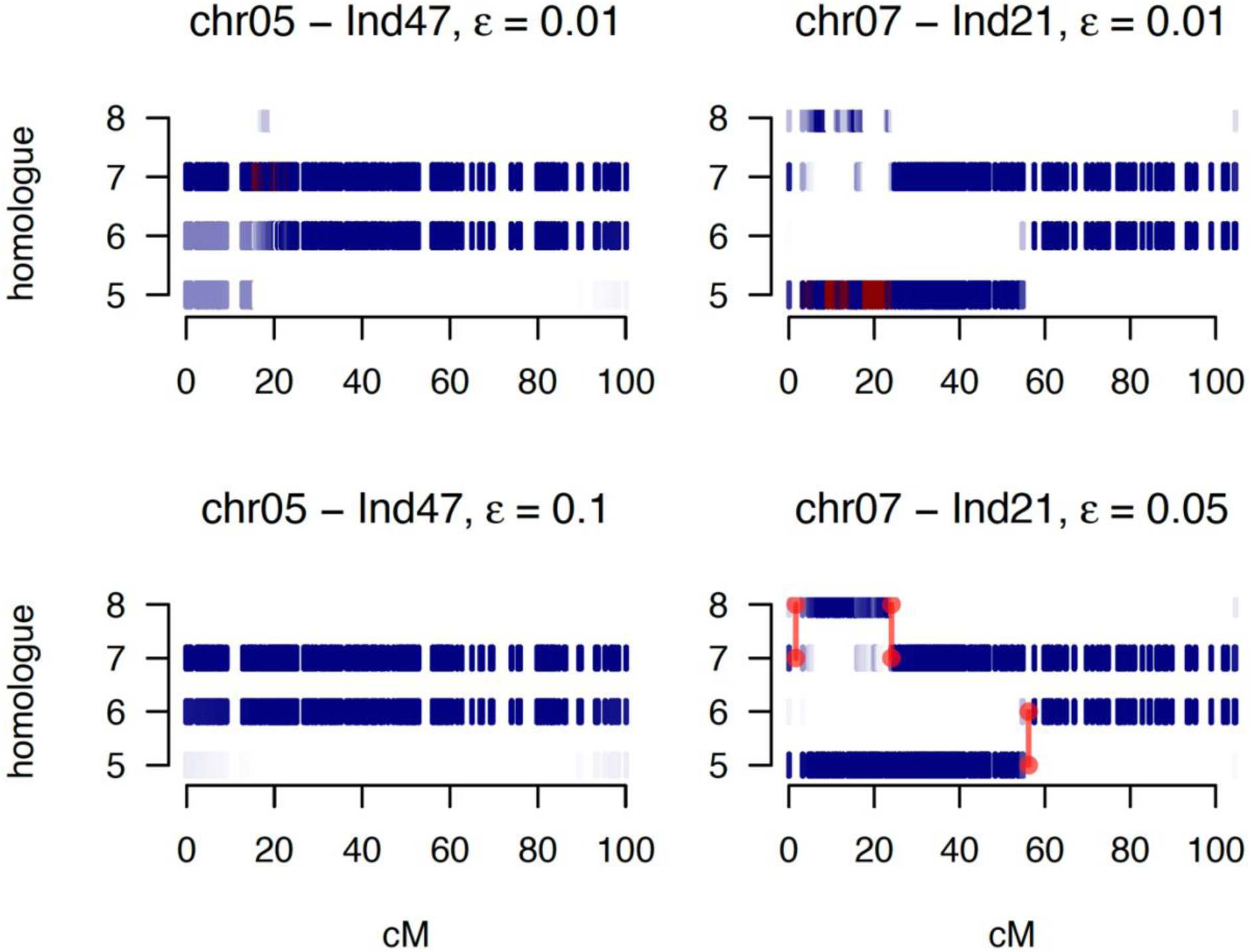
Visualization of posterior genotype (IBD) probabilities from a tetraploid Hidden Markov Model. Two individuals initially predicted to carry double reduction products (colored dark red segments, upper plots) at the default genotyping error prior of 0.01 were subsequently predicted to have arisen from bivalent pairing (within subgenomes) at error priors (0.1 and 0.05) that maximized the likelihood function. Homologue numbering 5 & 6 refers to phase 1 and 2 of LG5 (and LG7) while homologue 7 & 8 denote phase 1 and 2 of linkage group (LG) 15 (and LG17, respectively). Predicted recombination break-points are shown with red dots and lines in the bivalent-pairing case only.

**Figure S12:**
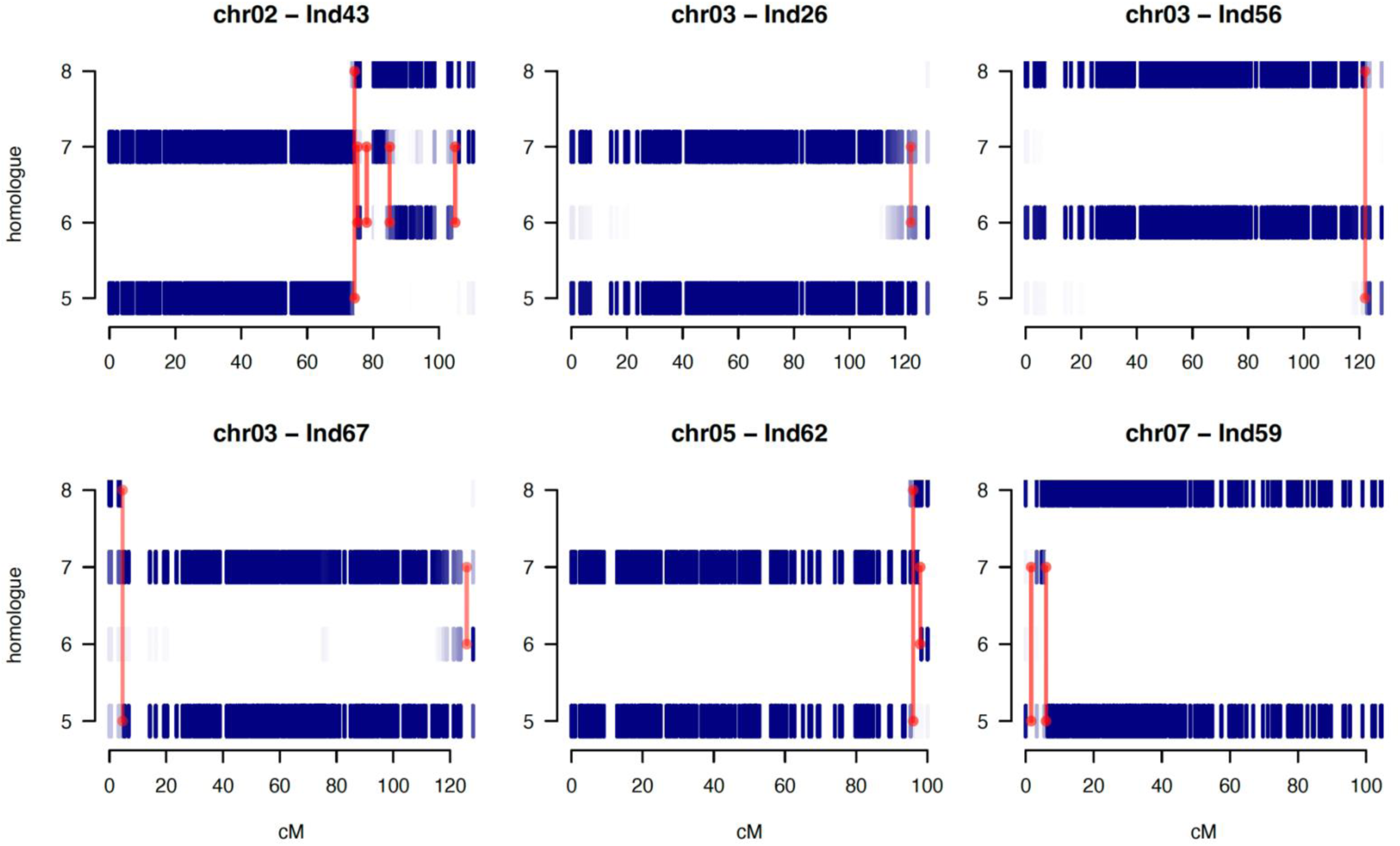
Visualization of posterior genotype (IBD) probabilities for individuals with predicted pairing and recombination across subgenomes (*i.e.* homoeologous exchanges). Allotetraploid pairing is only expected between homologues 5 & 6 or 7 & 8 (see legend of previous Figure S11 for further explanatory details).

